# Hepatic mitochondrial respiration is crucial for euthermia in complex III-deficient mice with impaired brown adipose tissue thermogenesis

**DOI:** 10.1101/2024.09.23.612616

**Authors:** Rishi Banerjee, Divya Upadhyay, Tomáš Zarybnický, Christa Kietz, Satu Kuure, Vineta Fellman, Janne Purhonen, Jukka Kallijärvi

**Affiliations:** Folkhälsan Research Center, Helsinki, Finland; Stem Cells and Metabolism Research Program, Faculty of Medicine, University of Helsinki, Finland; GM-unit, Laboratory Animal Center, Helsinki Institute of Life Science, University of Helsinki, Finland; Division of Clinical Microbiology, Department of Laboratory Medicine, Karolinska Institutet, Huddinge, Sweden; Children’s Hospital, University of Helsinki, Finland; Department of Clinical Sciences Lund, Pediatrics, Lund University, Sweden

**Keywords:** Mitochondrial disease, liver disease, gene therapy, respiratory complex III, adeno-associated virus, thermogenesis, UCP1, brown adipose tissue, hypothermia, sensory neuropathy

## Abstract

Liver is the key hub of systemic energy metabolism and growth regulation, yet its roles in mitochondrial disease pathophysiology remain relatively understudied. *Bcs1l^p.S78G^* knock-in mice, carrying a patient mutation causing respiratory complex III (CIII)-deficiency, present juvenile-onset liver and kidney disease, growth restriction, lipodystrophy, and early death. We restored CIII function in the hepatocytes of these mice using recombinant adeno-associated viral vectors (rAAVs) expressing wild-type *Bcs1l*. A single intraperitoneal injection of rAAVs into presymptomatic juvenile mice prevented liver disease, improved hypoglycemia and growth, normalized hepatic fuel utilization, and doubled the survival. The mutant mice showed hypothermia and brown adipose tissue (BAT) inflammation, and lacked BAT activation basally and upon acute cold challenge. Disrupted foot pad innervation suggested sensory neuropathy and impaired thermosensation as a contributor to the BAT inactivity. Surprisingly, the rAAV-treated mice maintained near-normal body temperature without significant effect on BAT. Increasing cellular respiration via transgenic alternative oxidase (AOX) was sufficient to prevent the hypothermia. The CIII-deficient mice did not reach euthermia until at an ambient temperature of 35°C, housing at which relieved metabolic stress and ameliorated hepatocyte senescence. We conclude that mitochondrial respiration in hepatocytes is essential for euthermia in mice. Our findings highlight the crucial role of the liver in thermoregulation, hypothermia as a consequence of mitochondrial dysfunction, and the therapeutic potential of rAAV-based gene delivery in a preclinical model of a multiorgan mitochondrial disease.

**Graphical abstract:** 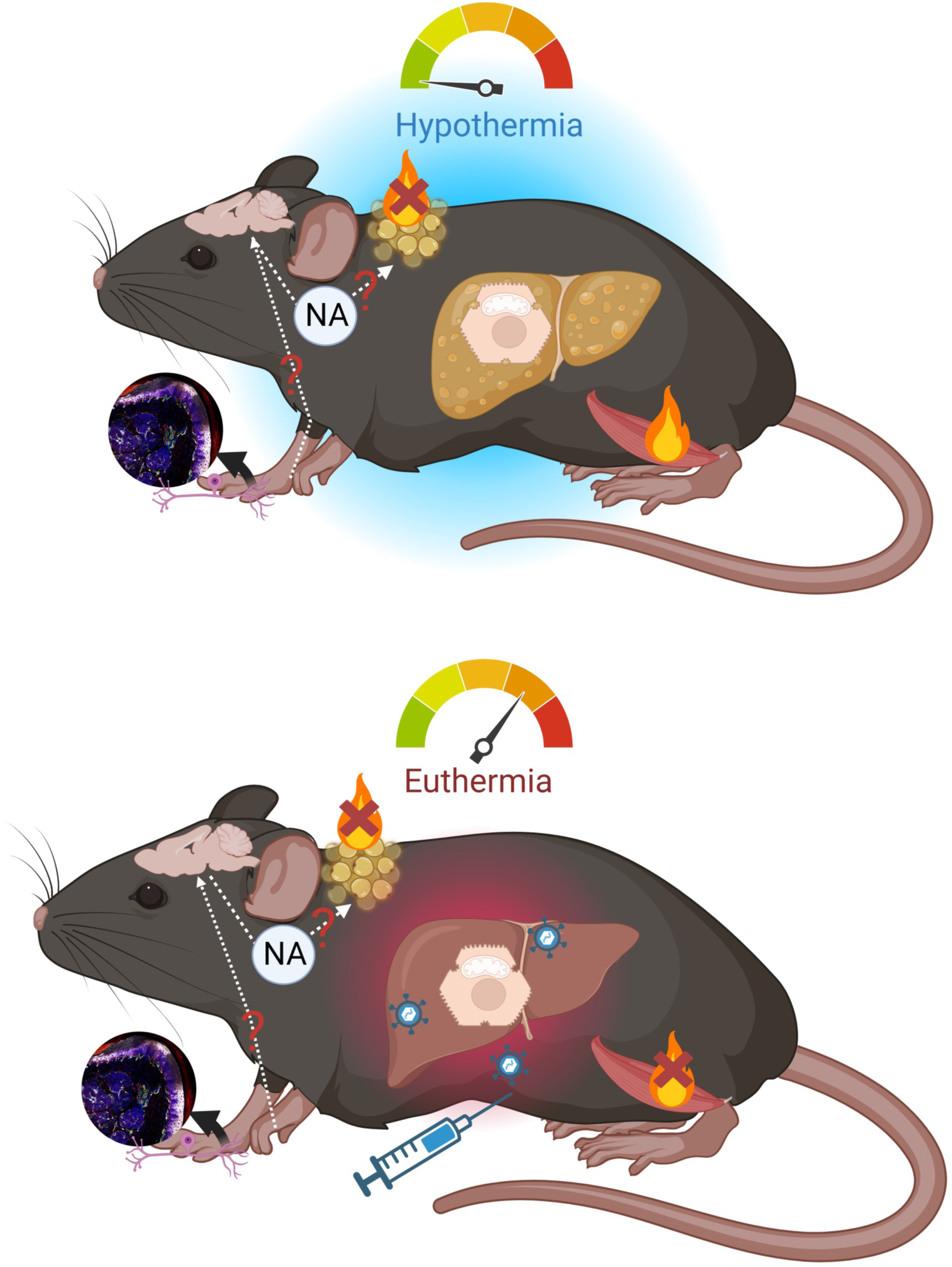

## Introduction

Mitochondria are cellular organelles that convert the chemical energy of nutrients into adenosinetriphosphate (ATP) and heat^1^. In mitochondrial diseases, the most common group of inherited errors of metabolisms, oxidative phosphorylation (OXPHOS) is directly or indirectly affected, potentially compromising both ATP and heat production^2^. Mitochondrial diseases can manifest in almost any tissue of the body, often present with ambiguous genotype-phenotype correlation and lack effective treatments^2^. Many aspects of the systemic energy metabolism and interorgan communication between the affected and non-affected tissues in mitochondrial diseases are poorly understood^3,4^. Complex III (CIII) deficiencies^5^ are a subgroup of mitochondrial diseases, among which the GRACILE syndrome (fetal-onset Growth Restriction, Aminoaciduria, Cholestasis, liver Iron overload, Lactic acidosis, and Early death during infancy) exhibits one of the most severe phenotypes^6,7^. *Bcs1l^p.S78G^* knock-in mice, carrying the GRACILE syndrome mutation, recapitulate most of the clinical manifestations^8–10^. In an mtDNA background (*mt-Cyb^p.D254N^*) that exacerbates the CIII deficiency, the homozygotes succumb to the metabolic crisis by 4-5 weeks of age^8,11^. The liver is one of the main affected organs in GRACILE syndrome and in the *Bcs1l^p.S78G^* mice. However, it is unknown to which degree the systemic metabolic phenotypes like growth restriction, lipodystrophy, and hypoglycemia are liver-dependent in this model^12^.

Recombinant adeno-associated viruses (rAAVs) are widely utilized delivery vectors of DNA in gene therapy strategies, partially due to their safety, being non-integrating viruses^13^. The liver is particularly amenable to viral transduction by rAAVs in mice^14^. To investigate the roles of the liver in the systemic energy metabolism in OXPHOS deficiency, we utilized rAAVs expressing wild-type mouse *Bcs1l* under a broad or a hepatocyte-specific^15^ promoter to restore CIII function and OXPHOS in the *Bcs1l^p.S78G^* mice. As outcomes, we assessed growth, liver disease progression, energy metabolism, body temperature, and survival. We show that the hepatocyte-specific rescue was sufficient to fully prevent the liver disease and ameliorate systemic manifestations, resulting in doubling of the survival. We discovered an essential role for basal liver thermogenesis in sustaining euthermia in the CIII-deficient mice. Finally, we show that forcing the hypothermic CIII-deficient mice to maintain normal body temperature via elevated housing temperature relieves their metabolic stress and hepatocyte senescence.

## Results

### Hepatocyte-targeted gene therapy doubles the survival of CIII-deficient mice

We injected presymptomatic (postnatal days, P19-23) mutant mice (genotype *Bcs1l^p.S78^*;*mt-Cyb^p.D254N^*, referred to as mutant (or MUT) from now on) with rAAV, assessed the disease progression and performed necropsy according to the timeline and experimental setup shown in Fig. 1A and B. Hepatocyte-specific ApoE enhancer and α1-antitrypsin (AAT) promoter^16^ drove mouse BCS1L or, for control, ehanced green fluorescent protein (EGFP) expression. The groups of mutant mice injected with rAAV carrying the *Bcs1l* or *EGFP* transgene will be referred to as AAT-*Bcs1l* and AAT-*EGFP,* respectively. Fluorescence microscopy of AAT-*EGFP* injected livers (Supplementary Fig. 1A) demonstrated high transduction efficiency, similar to previously published results^16^. The mutant livers that had received AAT-*Bcs1l* appeared visually as healthy as the WT livers one week after the injection (Supplementary Fig. 1B). qPCR analysis of total and virally-expressed *Bcs1l* at P28 confirmed the viral expression of *Bcs1l*, which resulted in an approximately 15-fold increased total *Bcs1l* mRNA in the liver (Fig. 1C).

**Figure 1.**
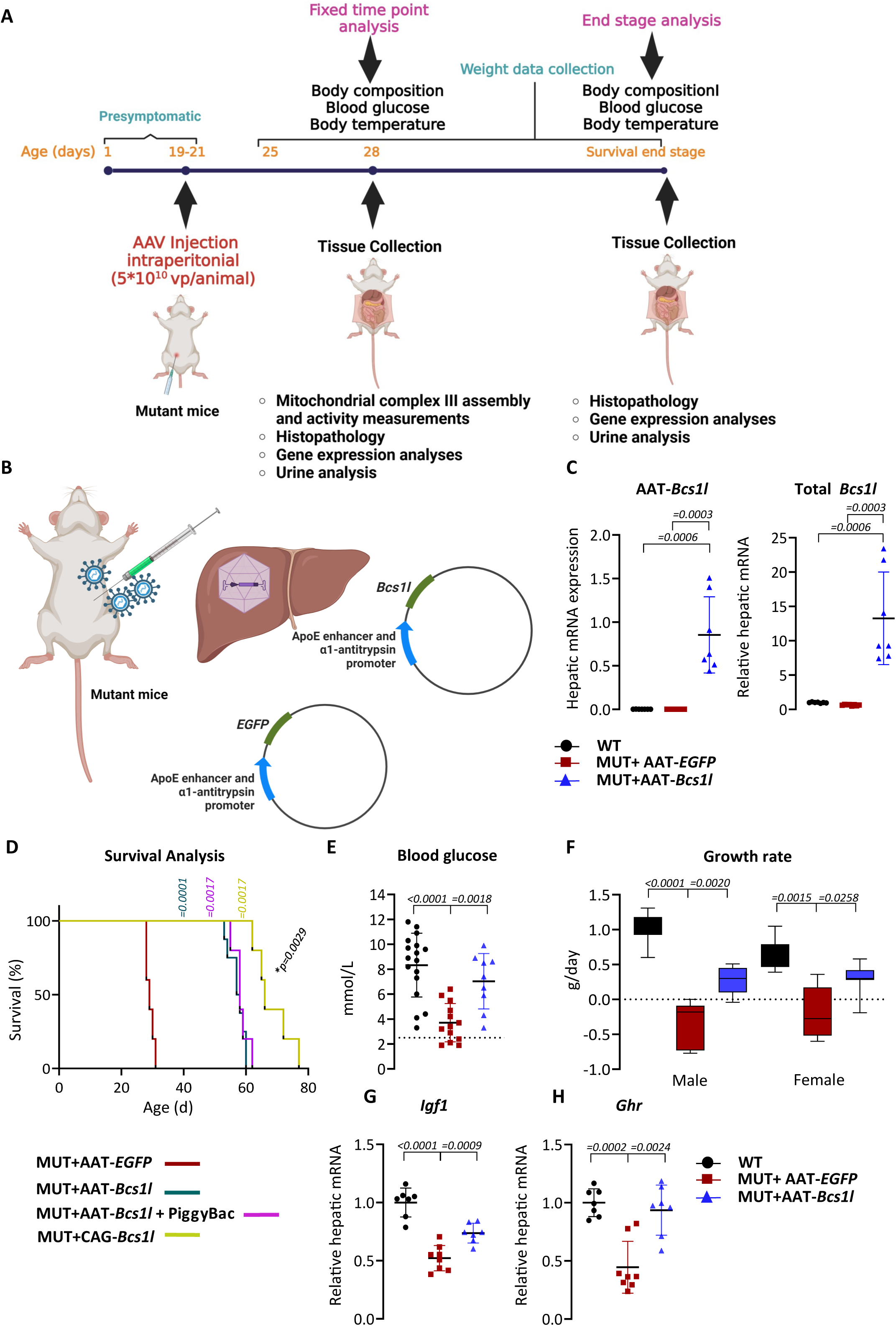
rAAV-based gene replacement therapy rescues the growth and doubles the survival of CIII-deficient mice. A) Schematic presentation of the experimental timeline and the time points of the investigations. B) Schematic presentation of the experimental setup. C) Virally-expressed and total *Bcs1l* mRNA at P28 (*n*=7-8/group). D) Survival curves of rAAV-*EGFP* and three different groups of mutant mice injected with different rAAV constructs (*n*=5-8/group). E) Blood glucose levels at P28 (*n*=9-17/group). The dotted line indicates the critical level of glucose (<2.5mmol/L) predicting spontaneous death. F) Sex segregated growth rate of the mice from P25 to P28 (n=6-13/group). G and H) Hepatic mRNA expression of insulin-like growth factor 1 (*Igf1*) and growth hormone receptor (*Ghr*) at P28 (*n*=7-8/group). Statistics: Mann-Whitney U test (C). One-way ANOVA followed by the selected pairwise comparisons with Welch’s t-statistics (E-H) and Survival data (D) were analyzed with log-rank test (Mantel-Cox). *p* values of the group comparisons are indicated above the panel. *\*p*, comparison between survival of AAT*-Bcs1l* and CAG*-Bcs1l* group. The error bars stand for standard deviation. All data points derive from independent mice.

At one month of age, the mutants injected with AAT-*Bcs1l* showed no signs of terminal deterioration, which consistently appeared in the AAT-*EGFP*-injected control group. Strikingly, the mean survival of the AAT-*Bcs1l*-injected mice was doubled to 58 days (Fig. 1D). *Bcs1l* expression was ∼2.4 fold higher compared to WTs at the end stage (Supplementary Fig. 1C). To ensure persistent expression in the growing liver, we co-injected PiggyBac transposase encoding rAAVs with AAT-*Bcs1l*. Inclusion of the transposase-expressing virus allowed genomic integration, which resulted in significantly higher end-stage hepatic *Bcs1l* expression (Supplementary Fig. 1C), but the survival did not extend further (Fig. 1D). This suggests that the eventual deterioration in the liver-targeted group was due to the disease progression in the other affected organs. To target also other tissues, we used a construct with the broadly active CAG promoter (CAG-*Bcs1l*). The broader *Bcs1l* expression did increase the survival further, albeit relatively modestly (15%), to a mean of 66 days (Fig. 1D). This vector led to a higher end-stage hepatic *Bcs1l* expression than the hepatocyte-specific promoter even without genomic intergration (Supplementary Fig. 1C). At P28, AAT-*Bcs1l* significantly improved the low blood glucose, and, importantly, none of the treated mice showed blood glucose less than <2.5mmol/L (Fig. 1E), thus avoiding lethal hypoglycemia^12,17,18^. However, at the end stage (P53-60), the treated mice again failed to maintain normal blood glucose (Supplementary Fig. 1E). While the control mutants started to lose weight after P25, the mutants treated with AAT-*Bcs1l* were able grow (Fig. 1F) and maintain hepatic insulin-like growth factor 1 (*Igf1*) and growth hormone receptor (*Ghr*) expression (Fig. 1G and H).

### Gene therapy restores hepatic CIII function and prevents hepatopathy

We isolated liver mitochondria and performed blue native gel electrophoresis (BNGE) and immunoblot analyses to assess CIII assembly. As shown previously^9^, the mutant mitochondria showed decreased fully-assembled CIII dimer (CIII_2_) based on the presence of UQCRFS1 (RISP). Almost all of the residual fully-assembled CIII_2_ was in the CI-CIII_2_ supercomplexes (SC), with free CIII_2_ being absent. The gene therapy efficiently increased the levels of fully-assembled CIII_2_ (Fig. 2A). In isolated mutant liver mitochondria, the mean CIII activity was around 26% of WT. The gene therapy increased the CIII activity to approximately 74% of WT (Fig. 2B) and improved hepatic ATP levels (Fig. 2C). The increased mitochondrial biogenesis marker *Ppargc1a* (*Pgc-1α*) (Fig. 2D) and mitochondrial mass markers VDAC1 and HSP60 returned to WT level in the AAT-*Bcs1l*-injected mice (Fig. 2E). HSP60 immunostaining of liver tissue sections showed an abnormal clustered pattern of mitochondria in a fraction of mutant hepatocytes, which was again fully prevented by the AAT-*Bcs1l* (Supplementary Fig. 1F).

**Figure 2.**
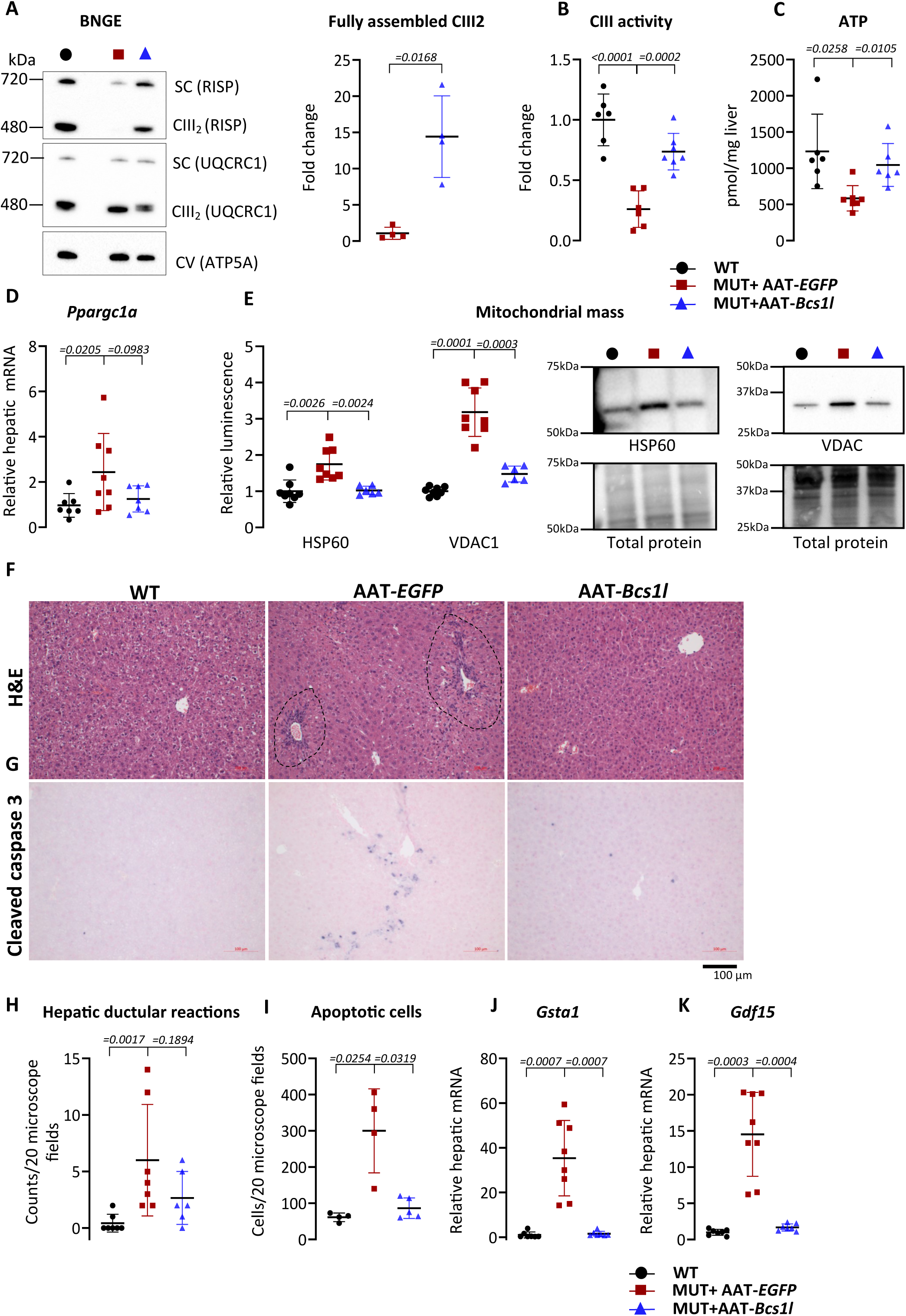
Hepatocyte-targeted gene replacement corrects hepatic CIII assembly and prevents hepatopathy. A) Representative blue-native PAGE blot of UQCRFS1 (RISP) and UQCRC1 assembled into free CIII_2_ and supercomplexes (SCs) in isolated liver mitochondria at P28, and RISP per total UQCRC1 ratio (*n*=4/group). B) CIII activity normalized by total protein in liver isolated mitochondria at P28 (*n* = 4/group). C) ATP in liver at P28 (*n* = 6-7/group). D) mRNA expression of a transcriptional regulator of mitochondrial biogenesis *Ppargc1a (PGC-1α)*, from P28 liver (*n*=7-8/group). E) Western blot quantification and representative blots of HSP60 and VDAC1 as markers of mitochondrial mass from P28 liver lysates (*n*=6-8/group). F and G) Representative images of H&E-stained liver sections (*n*=6-7/group), showing tissue morphology and expansion of portal areas (indicated by dotted lines), and liver sections immunostained for apoptotic cell marker, cleaved caspase 3 (cc3) (*n*=5/group) at P28. H and I) Quantification of hepatic ductular reactions and the apoptotic cells from H&E-stained and cc3-immunostained sections, respectively. J&K) *Gsta1* and *Gdf15* mRNA expression from P28 liver (*n*=7-8/group). Statistics: Welch’s *t*-test (A), one-way ANOVA followed by the selected pairwise comparisons with Welch’s t-statistics (B-E&I-K) and Mann-Whitney U test (H). P values of the group comparisons are indicated above the panel. The error bars stand for standard deviation. All data points derive from independent mice.

Histopathological analysis of the P28 mutant mice showed early-stage hepatopathy characterized by expansion of portal areas, increased ductular reactions and cell death (Fig. 2F-I). The AAT-*Bcs1l* fully prevented these changes (Fig. 2F-I), as well as the upregulation of the disease marker gene^19^ glutathione S-transferase 1 (*Gsta1*) at P28 (Fig 2J) and the mitochondrial dysfunction-associated mitokine, growth-differentiation factor 15 (*Gdf15*) at P28 (Fig. 2K). The hepatic ductular reactions and *Gdf15* expression were increased at the end stage (Supplementary Fig. 2A-F), indicating eventual liver pathology despite the remaining WT *Bcs1l* expression.

To assess possible liver-dependent systemic effects in other organs, we analyzed kidney histology. Proximal tubulopathy, presumably due to damage to the mitochondria-rich tubular epithelial cells, is another prominent visceral phenotype in GRACILE syndrome and in the *Bcs1l^p.S78^* mice^6,8,9^. This manifests as increased cell death and loss of cortex volume already at P28 (Supplementary Fig. 3A, B, E anf F). Surprisingly, despite the fact that the number of apoptotic cells was not decreased (Supplementary Fig. 3B and F), the kidney cortex thickness was slightly increased by AAT-*Bcs1l* (Supplementary Fig. 3A and E), suggesting a systemic effect from the rescued liver. The treated mice showed normal kidney cortex thickness even at the end-stage (Supplementary Fig. 3C, D and G-J).

### Hepatocyte-targeted gene therapy reverses hypothermia in CIII-deficient mice

We found that the symptomatic (>P25) mutant mice had hypothermic body temperature ranging from below 32 to 36°C at P28-30 (Fig. 3B). Cold-inducible genes RNA-binding motif protein 3 (*Rbm3*) and cold-inducible RNA-binding protein variant 4 (*Cirbp4*) were upregulated in the liver (Supplementary Fig. 4A), indicating a transcriptional response to hypothermia Surprisingly, the body temperature was normalized to near WT level by AAT-*Bcs1l* (Fig. 3A and B), indicating that the hypothermia was a consequence of the loss of CIII function in hepatocytes. Having discovered that restoring CIII function in hepatocytes reversed the hypothermia, we expressed alternative oxidase (AOX), a non-mammalian mitochondrial enzymes that can restore coenzyme Q (CoQ) oxidation but not OXPHOS when the CIII-CIV segment is blocked. It can increase heat generation indirectly by decreasing OXPHOS efficiency and directly by reducing molecular oxygen to water^20^. Hepatocyte-specific rAAV expression of *Crassostrea gigas* AOX (AAT-*CgAOX*) had no effect on the hypothermia (Supplementary Fig. 4B) or hepatic cold inducible gene expression (Supplementary Fig. 4C). In contrast, mutant mice with broad transgenic expression of *Ciona intestinalis* AOX, which we have shown to prevent liver disease and multiple other tissue pathology in the *Bcs1l^p.S78^*mice^9,10^, maintained near-normal core temperature (Fig. 3B). These data corroborate that cellular respiration in hepatocytes was crucial for the liver to maintain body temperature.

**Figure 3.**
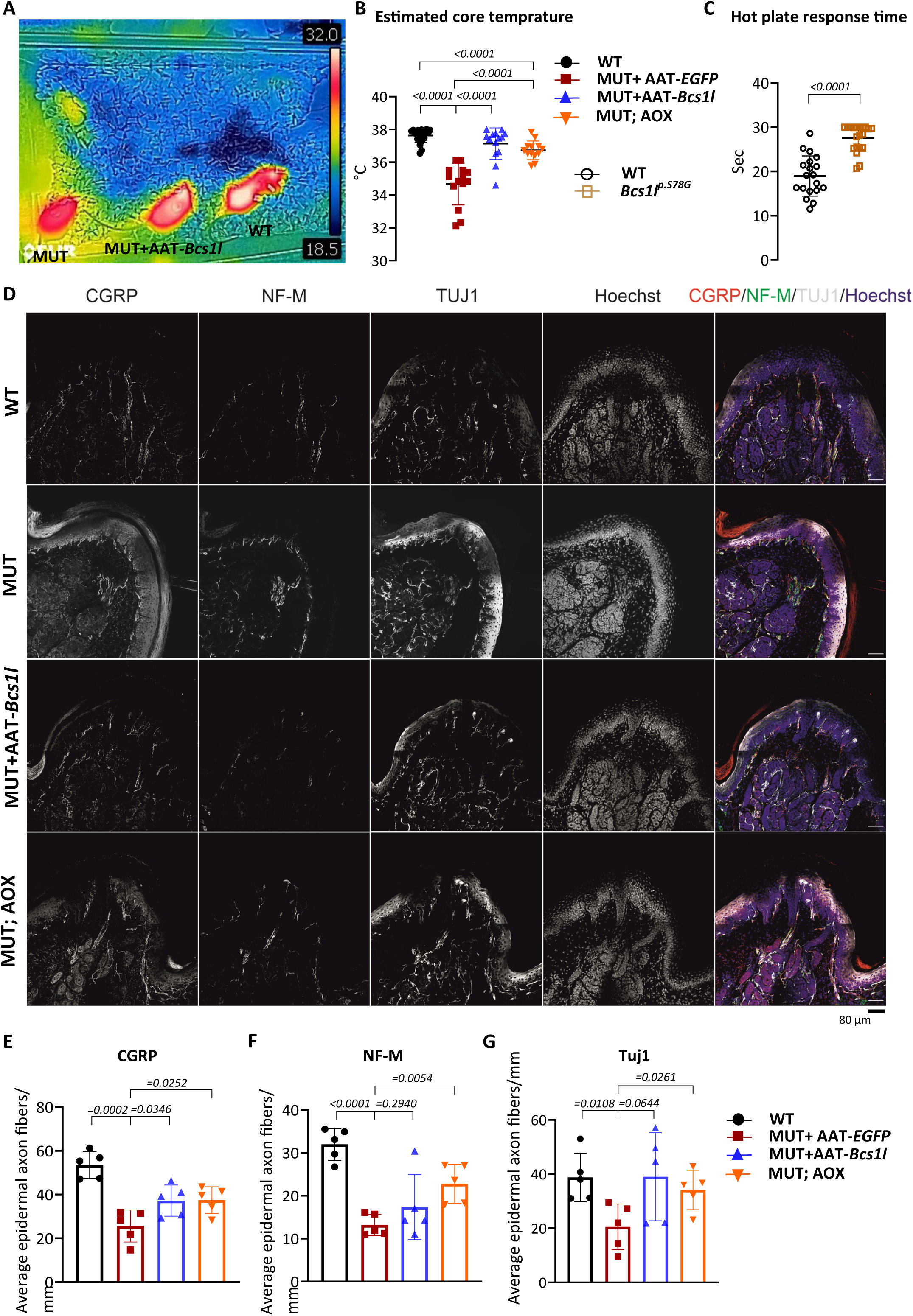
Hypothermia and impaired thermosensation in CIII-deficient mice. A) Representative infrared image of P28 mice. B) Estimated core temperature of P28 mice (n=13-21/group). C) Hot plate response time of P100 mice (WT mtDNA background) (n=19-20/group). D-G) Representative images of WT and mutant foot pad neurons stained with CGRP, NF-M and TUJ1, and the quantification of average epidermal axon fibers (*n*=5/group). Statistics: one-way ANOVA followed by the selected pairwise comparisons with Welch’s t-statistics (B, C & E-G). The error bars stand for standard deviation. All data points derive from independent mice.

### CIII-deficient mice show disrupted sensory innervation of the skin

The inability of the CIII-deficient mice to maintain body temperature could be due to defective thermosensation, thermogenesis, thermoregulation, or all of these. Thermoregulation is based on the sensory nervous system that detects the ambient and body temperatures via cold- and heat-sensitive ion channels in sympathetic nerve termini throughout the body^21^. To assess this, we inspected hot plate assay data from adult (P100) *Bcs1l^p.S78G^* mice^9^. The data showed significantly increased response times in the mutant mice (Fig. 3C), which suggests compromised thermosensation and sensory neuropathy. Immunofluorescent labelling of peripheral nerves in the hind limb foot pads showed that the mutant foot pads exhibited fragmented β3 tubulin (TUJ1, pan-neuronal marker) labelling and minimal calcitonin gene-related peptide (CGRP, sensory neuron marker) and neurofilament M (NF-M) signals at the epidermal/dermal boundary of the foot pad (Fig. 3D-G). Quantification of epidermal axon fibers showed that the *AAT-Bcs1l* treatment increased the number of CGRP-positive axons in the mutant footpad epidermis (Fig. 3D-G), but unlike AOX transgene, had minimal effect on the NF-M and TUJ1-positive neuronal subpopulations (Fig. 3D-G). These data demonstrate structurally deficient skin sensory innervation in the *Bcs1l^p.S78G^*mice, which may affect thermosensation and subsequent adaptive thermogenesis by BAT and skeletal muscle. The data also suggest that the gene therapy and AOX expression had a subtle correcting effect on the skin sensory innervation.

### BAT is unresponsive to cold in CIII-deficient mice

BAT depots undergo thermogenesis-promoting functional remodeling and gene expression changes, such as mitochondrial biogenesis and UCP1 induction, in response to neuronal cold signals coming from the hypothalamus, and to circulatory catecholamines^22^. To assess if the histologically apparent peripheral neuropathy affected BAT stimulation, we collected and analysed interscapular BAT (iBAT) depots from the AAT-*Bcs1l*-injected and control mice. The mass of the iBAT depots was not significantly affected by genotype or treatment (Fig. 4A). Unexpectedly, markers of mitochondrial mass (SDHA/CII and UQCRC1/CIII) were unaffected in the mutant BAT (Fig. 4B). However, iBAT mitochondrial mass was increased in the transgenic AOX-expressing mice (Fig. 4B). In the mutant iBAT, the ratio of CIII subunits UQCRFS1 (RISP) and UQCRC2 was decreased by 40% (Fig. 4C), and CIII activity was approximately 35% of WT (Fig. 4D). The AAT-*Bcs1l* injection did not affect the the ratio of CIII subunits and CIII activity in iBAT, as expected (Fig. 4C and D).

**Figure 4.**
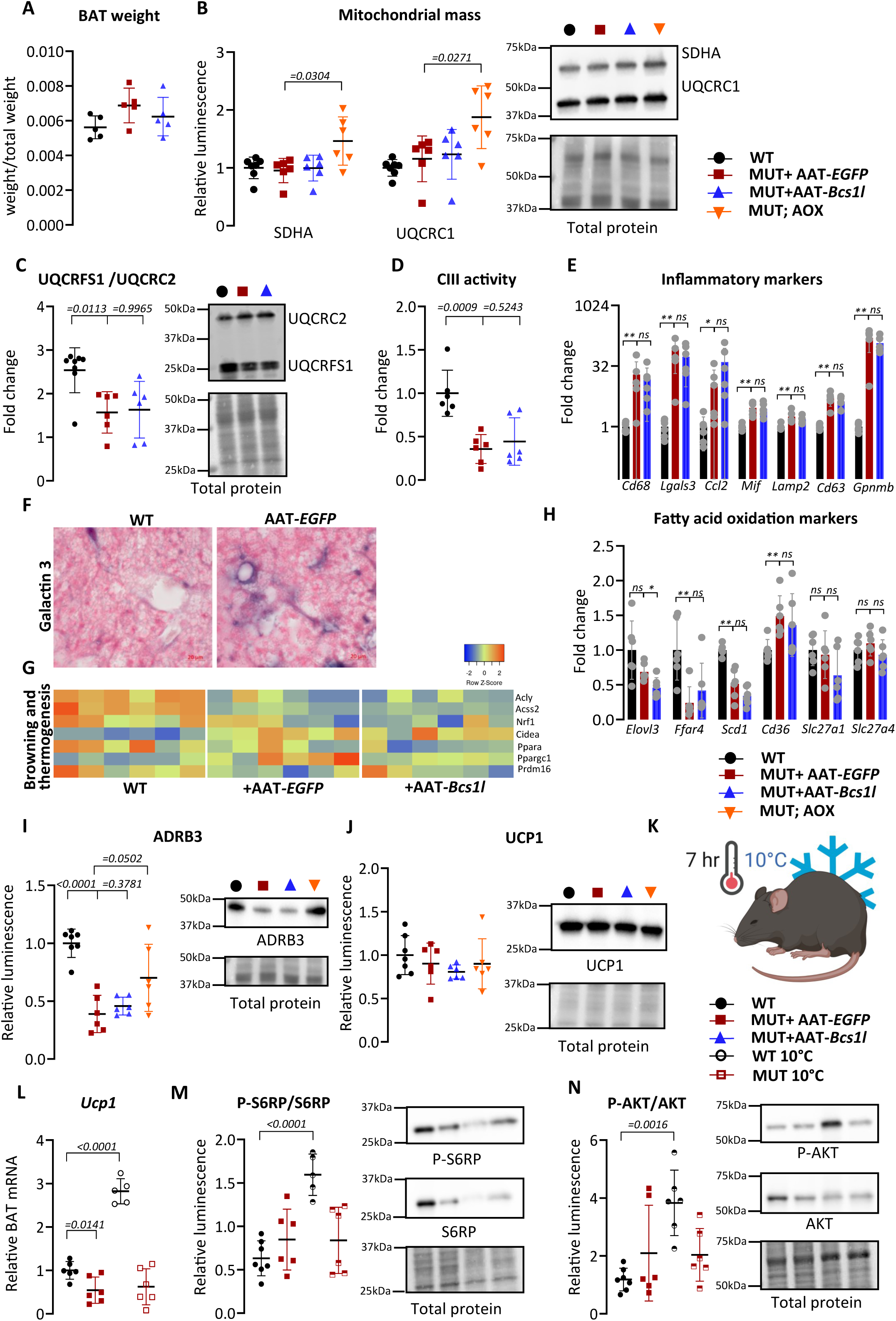
Impaired brown adipose tissue thermoregulation in mutant mice. A) Total BAT tissue weight at P28, normalized by weight of the animal. B and C) Western blot quantification and representative blot of SDHA, UQCRC1, UQCRFS1 and UQCRC2 from P28 BAT lysates (*n*=6/group). D) CIII activity in P28 BAT normalized by CIV activity (*n* = 6/group). E) Differentially expressed genes in BAT transcriptome related to inflammation. F) Representative images of BAT sections immunostained for inflammatory marker, Galactin 3 (*n*=4/group) at P28. G) Heat map visualization of gene expression related to browning and thermogenesis in BAT (n=6/group) H) Differentially expressed genes in BAT transcriptome related to fatty acid oxidation. I and J) Western blot quantification and representative blot of adrenergic receptor (ADRB3) and uncoupling protein 1 (UCP1) from P28 BAT lysates (*n*=6/group). K) Schematic presentation of the experimental setup of cold treatment. L) *Ucp1* mRNA expression from P28 BAT (n=6/group). M and N) Western blot quantification of the Ribosomal protein S6-Ser240/244 (S6RP) and Akt-Ser473 (AKT) phosphorylation from P28 BAT lysates (*n*=6/group). Statistics: one-way ANOVA followed by the selected pairwise comparisons with Welch’s t-statistics (A-D, H-J and L-N) and Mann-Whitney U test (E). *\*\*p ≤ 0.01, *p ≤ 0.05.* The error bars stand for standard deviation. All data points derive from independent mice.

The low CIII activity alluded that BAT-intrinsic mechanisms dependent on mitochondrial respiration may contribute to the BAT inactivity. iBAT gene expression analysed showed upregulation of the BAT inflammation- and macrophage-related genes^23,24^ *Cd68*, *Lgals3*, *Ccl2, Mif* and *Lamp2*, and AAT-*Bcs1l* did not correct these changes (Fig. 4E). Interestingly, markers for a specific fibrogenic population of macrophages (Fab5, CD9^+^TREM2^+^ and expressing SPP1, GPNMB, FABP5, and CD63)^25^ were highly increased in the liver, and this was fully prevented by AAT-*Bcs1l* (Supplementary Fig. 5A). The Fab5-related gene expression data was less robust for iBAT, but *Cd63* and *Gpnmb* were highly upregulated (Fig. Fig. 4E). In line with that, immunostaining showed a highly increased number of Galectin-3 (also called MAC-2)-positive cells (“crown-like structures”) in the mutant iBAT (Fig. 4F). Adipocyte browning and thermogenesis-related gene expression was suppressed in the mutant mice, and this was not corrected by AAT-*Bcs1l* (Fig. 4G). Moreover, expression of key fatty acid metabolism genes (*Elovl3, Ffar4, Scd1, Cd36, Slc27a1, Slc27a4*) was largely decreased or unchanged in the mutant iBAT (Fig. 4H), in contrast to the upregulation typically seen in active BAT^26,27^. At global gene expression level, the hepatocyte-specific restoration of CIII function had a minimal effect on iBAT, as the Venn diagram shows 187 overlapping genes in WT vs. AAT-*Bcs1l* iBAT but 520 overlapping genes AAT-EGFP vs. AAT-*Bcs1l* iBAT (Supplementary Fig. 5B).

Unlike the neuronal markers in foot pads, tyrosine hydroxylase (TH), a marker for sympathetic nerve terminals, was not decreased in the mutant iBAT (Supplementary Fig. 5C). When activated by hypothalamic signals, these nerves release noradrenaline (NA), which binds to β3 adrenergic receptors (ADRB3) on adipocytes, stimulating BAT thermogenesis^22,28^. Both ADRB3 protein and *Adrb3* mRNA were downregulated (by 60% and 90%, respectively) in the mutant iBAT (Fig. 4I and Supplementary Fig. 5D), which suggests decreased adrenergic signaling to iBAT despite the hypothermia. Persistent adrenergic stimulation has been shown to downregulate ADRB3 via TRIB1 (tribbles pseudokinase 1) upregulation^29^. However, *Trib1* was slightly downregulated in the mutant iBAT (Supplementary Fig. 5E), speaking against this mechanism. *Adrb3* remained suppressed in the AAT-*Bcs1l*-injected mice (Fig. 4I and Supplementary Fig. 5D) Taken together, the normalization of the core temperature by AAT-*Bcs1l* did not depend on the BAT. In contrast, the transgenic AOX-expressing iBAT showed partial normalization of ADRB3 levels (Fig. 4I and Supplementary Fig. 5D).

Dissipitation of the mitochondrial membrane potential via uncoupling protein 1 (UCP1) is the main heat-generating mechanism in BAT. Upon acute cold exposure, ADRB3 signaling rapidly activates UCP1 transcription^30^. Despite their inherent hypothermia, iBAT *Ucp1* mRNA was slightly downregulated in mutant iBAT, indicating lack of transcriptional response to cold, and this was not effected by AAT-*Bcs1l* or by transgenic AOX expression. (Supplementary Fig. 5F). UCP1 protein levels were unchanged in all groups of mutant mice (Fig. 4J) housed at room temperature (RT). To examine the responsiveness of BAT to an acute cold exposure, we kept the mice at 10°C for 7 hrs (Fig. 4K and Supplementary Fig 5G). This acute cold exposure caused an expected approximately 3-fold induction of *Ucp1* mRNA in WT iBAT (Fig. 4L), even though the UCP1 protein level did not change (Supplementary Fig 5H). Strikingly, *Ucp1* mRNA was not induced at all in the mutant iBAT (Fig. 4L). Cold-induced ADRB3 receptor activation is followed by protein kinase A and mTOR signalling to upregulate UCP1 expression^28,31^. Indeed, upon the acute cold exposure both Akt Ser473 and S6 ribosomal protein Ser240/244 phosphorylation were increased in WT mice, but not in the mutant mice (Fig. 4M and N). These data confirmed that the iBAT of the mutant mice was unresponsive to both internal and ambient cold temperature. Taken together, we conclude that the low CIII activity in BAT, possibly in combination with decreased sympathetic stimulation, lead to BAT inflammation and dysfunction, contributing to the compromised BAT thermogenesis in the CIII-deficient mice.

### Restoration of CIII function in hepatocytes reverses a thermogenic shift in skeletal muscle

Skeletal muscle is the largest (45-55% of body mass) adaptive thermogenic organ in animals, and skeletal muscle thermogenesis can compensate the loss of BAT thermogenesis in mice^32^. Skeletal muscle cells use both myosin-mediated ATP hydrolysis and Ca^2+^ transport-driven ATP hydrolysis by the SERCA pump for heat generation. Sarcolipin (SLN)-mediated uncoupling of the SERCA Ca^2+^ pump is the key non-shivering thermogenesis mechanism in the skeletal muscle^33^. Increased expression of *Sln, Ryr1* (encoding Ryanodine receptor 1, the sarcoplasmic reticulum calcium channel) and *Adrb2* (β2 adrenergic receptor) in the quadriceps (Fig. 5A) suggested that thermogenic signaling to skeletal muscle was functional and activated. To aid thermogenesis, SLN promotes oxidative energy metabolism via *Ppargc1a (Pgc-1α)*^34,35^, which was indeed induced in the mutant skeletal muscle (Fig. 5B). Upon decreased glucose availability, such as during prolonged exercise, skeletal muscle upregulates fatty acid oxidation^36^. Expression of pyruvate dehydrogenase kinase 4 (*Pdk4*), which inactivates the pyruvate dehydrogenase complex to dampen glucose oxidation, and fatty acid transporter *Cd36* were robustly upregulated in the mutant muscle (Fig. 5B) indicating a metabolic shift from gluocse to fatty acid utilization (Fig. 5C). Strikingly, the upregulation of *Pdk4* was fully prevented and *Cd36* normalized by AAT-*Bcs1l* (Fig. 5B), indicating a stong metabolic crosstalk between the liver and skeletal muscle. Also global gene expression analyses showed a significant systemic effect on skeletal muscle gene expression by the hepatocyte-targeted gene therapy (Fig. 5D). Taken together, the skeletal muscle data showed that the CIII-deficient mice were apparently able to sense hypothermia to activate skeletal muscle non-shivering thermogenesis, but this was insufficient in the non-treated mutant mice and unnecessary when CIII activity was restored in the liver.

**Figure 5.**
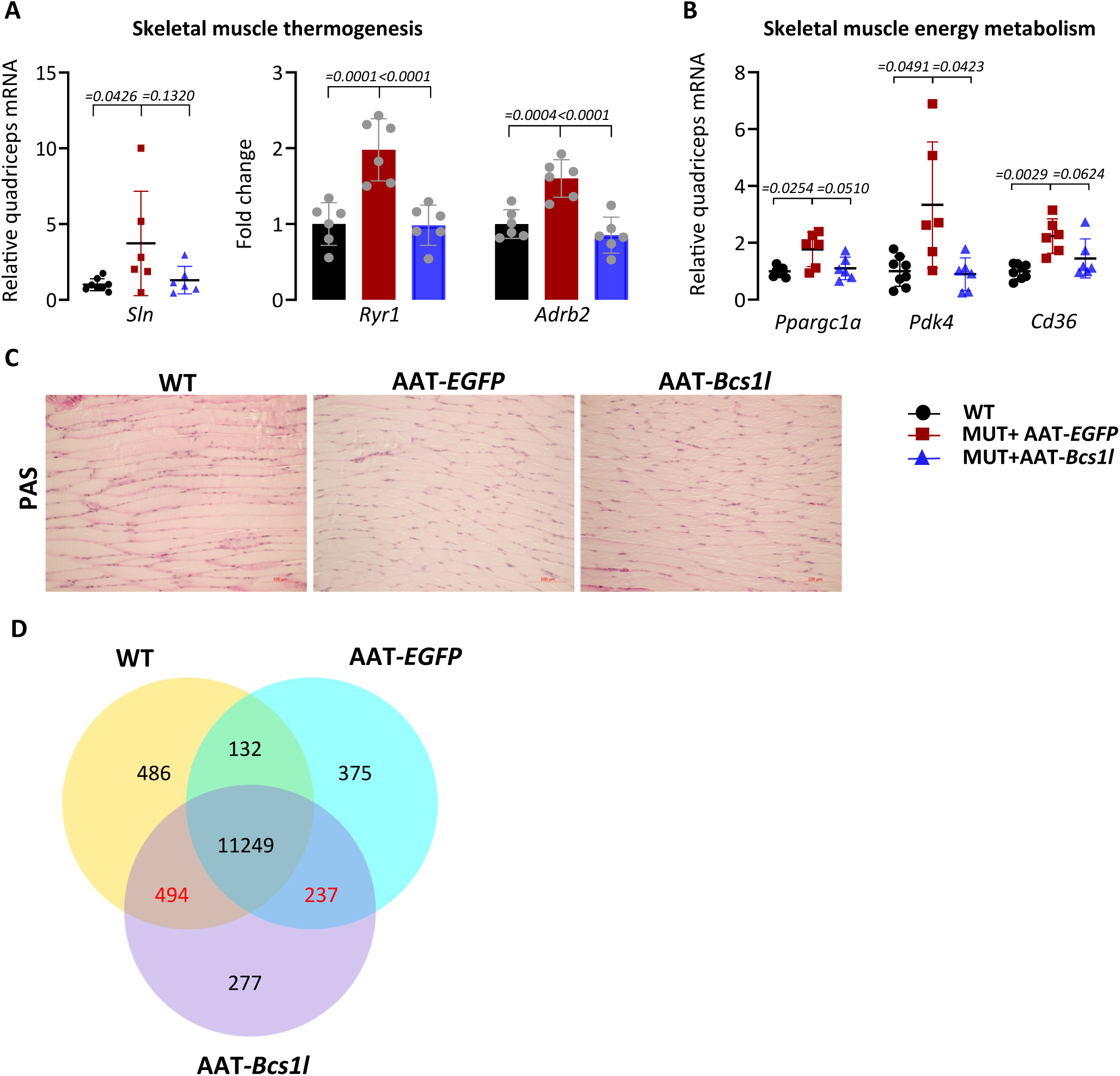
Effect of liver-targeted gene replacement on skeletal muscle thermogenesis. A) mRNA expression of *Sln, Ryr1*, and *Adrb2* from P28 quadriceps (n=6-8/group). B) *Ppargc1*, *Pdk4* and *Cd36* mRNA from P28 quadriceps (n=6-8/group). C) Representative images of periodic acid-Schiff (PAS)-stained quadriceps sections (n=5/group), detecting glycogen at P28. D) Differentially expressed genes in skeletal muscle transcriptome. Statistics: one-way ANOVA followed by the selected pairwise comparisons with Welch’s t-statistics. The error bars stand for standard deviation. All data points derive from independent mice.

### Restoration basal liver metabolism sustains euthermia in CIII-deficient mice

rAAV-mediated restoration of CIII function in hepatocytes was sufficient to fully prevent hypothermia and this effect was not mediated by skeletal muscle or BAT thermogenesis, as we show above. UCP1 is not expressed in the liver, precluding adaptive heat generation by mitochondrial uncoupling. Thus, the hepatic heat originates from basal mitochondrial and non-mitochondrial metabolisms. Hepatocytes preferentially use fatty acids as their primary energy source even under basal conditions. The low blood glucose (Fig. 1E) and glycogen depletion (Fig. 6A and B) in the CIII-deficient mice forces a transcriptional switch to boost fatty acid oxidation (FAO) even further. Marking this, Both *Pdk4* and *Cd36* were highly induced in the mutant liver (Fig. 6C, D). Despite increased fatty acid flux to the liver, which we have shown before^12^, the expression of *Pparα*, a major driver of fatty acid oxidation was significantly decreased (Fig 6E). Global gene expression analyses suggested an overall repression of FAO (Fig 6F), particularly mitochondrial β-oxidation (Fig 6G), which is directly linked to CIII function via the mitochondrial coenzyme Q pool. This was reflected in the typical microvesicular fat accumulation in the mutant liver (Fig. 6A), compatible with insufficient FAO capacity. AAT-*Bcs1l* largely prevented the fat accumulation (Fig. 6A), glycogen depletion (Fig. 6B), and the transcriptional changes (Fig. 6C-E), indicating normalization of hepatic energy metabolism, as expected. Accordingly, phosphorylation of AMP-dependent protein kinase (AMPK), a cellular sensor of ATP and glucose availability, was normalized to WT level (Fig 6H). As an additional test for the critical role of basal metabolic heat, we injected the mutant mice with thyroid horme (T3, triiodothyronine) on four consecutive days before sample collection at P28. Thyroid hormone strongly stimulates basal metabolism, thereby increasing non-adaptive thermogenesis in healthy normal mice^37,38^. T3 did not increase the body temperature of the mutant mice but further decreased it, indicating that they had no capacity to increase basal metabolism (Supplementary Fig. 6A). In summary, restoring mitochondrial respiration in hepatocytes reestablished hepatic basal metabolism and heat production (Fig 6I), which was sufficient to sustain euthermia despite the persisting CIII deficiency in other tissues including the skeletal muscle and BAT.

**Figure 6.**
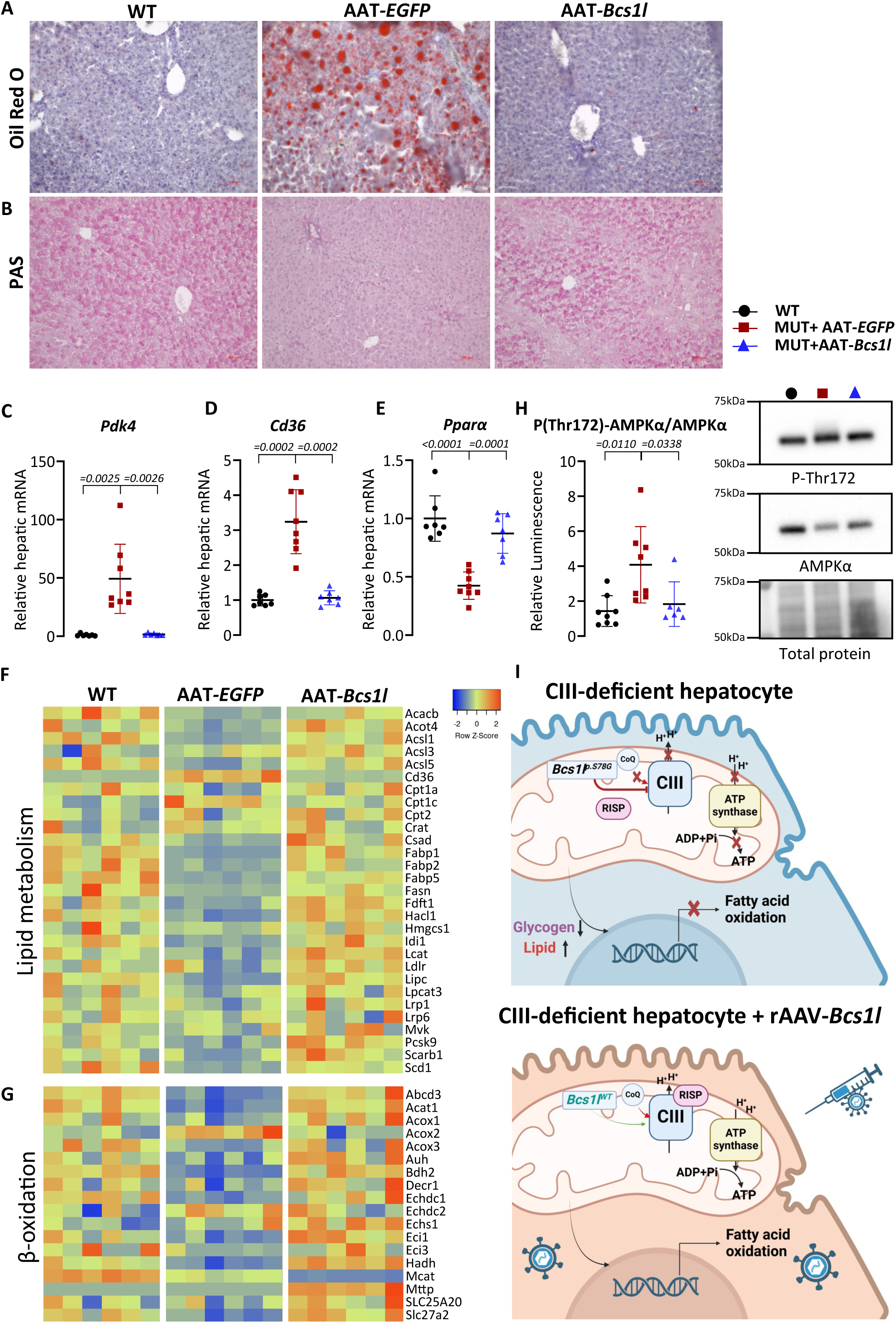
Restoration of CIII function in hepatocytes re-establishes the basal liver metabolism. A&B) Representative images of Oil-Red-O staining of liver cryosections (n=4/group) showing lipid accumulation and periodic acid-Schiff (PAS) stained liver sections (n=5/group) detecting glycogen at P28. C-E) *Pdk4, Cd36*, and *Pparα* mRNA expression from P28 liver (*n*=7-8/group). F&G) Heat map visualization of gene expression related to lipid metabolism and β-oxidation in liver (n=6/group) H) Western blot quantification of the phosphorylation status of AMPKα from P28 liver lysates (*n*=6-8/group) I) Schematic presentation of the of the metabolic rewiring of hepatocytes followed by AAT-*Bcs1l* injection Statistics: one-way ANOVA followed by the selected pairwise comparisons with Welch’s t-statistics (C-F). The error bars stand for standard deviation. All data points derive from independent mice.

### Housing at 35°C relieves metabolic stress and prevents hepatocyte senescence in CIII-deficient mice

Tissue growth and repair require considerable amounts of energy and biosynthetic resources. The *Bcs1l^p.S78G^*mice show regenerative hepatocyte proliferation, which in the face of their depleted nucleotide pools and other biosynthetic resources leads to DNA damage, cell cycle arrest and senescence^10^. Restoration of CIII function in hepatocytes via AAT-*Bcs1l* decreased the DNA damage/senescence marker γH2AX (Fig. 7A) and abolished the cell cycle arrest marker CDKN1A (p21) (Fig. 7B). It also decreased the cell proliferation markers PCNA and cyclin A2 to wild-type level (Supplementary Fig. 6B), indicating prevention of liver damage-induced hepatocyte regeneration. To probe the effects of the hypothermia-related metabolic stress on tissue regeneration, we first housed the mice at 30°C, a typical thermoneutral temperature for WT mice. Surprisingly, the *Bcs1l^p.S78G^* mice were not able to reach normal body temperature at 30°C, which suggested that their thermoneutral temperature was higher (Supplementary Fig. 6C). We next housed mutant mice at 35°C from P25, the age at which we estimated the hypothermia to begin, until P28 (Fig. 7C). At this temperature the mutant mice showed no typical shivering or decreased mobility and were able to maintain the core temperature at WT level (Fig. 7D), despite that their blood glucose was not improved (Fig. 7E). Hepatic AMP-dependent protein kinases (AMPK) phosphorylation was significantly decreased at 35°C in both mutant and WT mice (Fig 7F), indicating improved energy homeostasis by decreased energy expenditure for thermogenesis. Also the upregulation of the liver disease marker gene^19^ glutathione S-transferase 1 (*Gsta1*) was dampened (Fig 7G). In contrast, *Gdf15*, a marker for mitochondrial stress, was not decreased, underscoring that the mitochondrial stress due to the CIII deficiency was not alleviated (Fig. 7H). The forced euthermia completely prevented the the upregulation of the senescence markers γH2AX and CDKN1A/p21 (Fig. 7I and J). Taken together, these data show that inadequate thermogenesis is a crucial component of the pathophysiology in CIII-deficient mice and that hepatic mitochondrial respiration is necessary for euthermia in juvenile mice.

**Figure 7.**
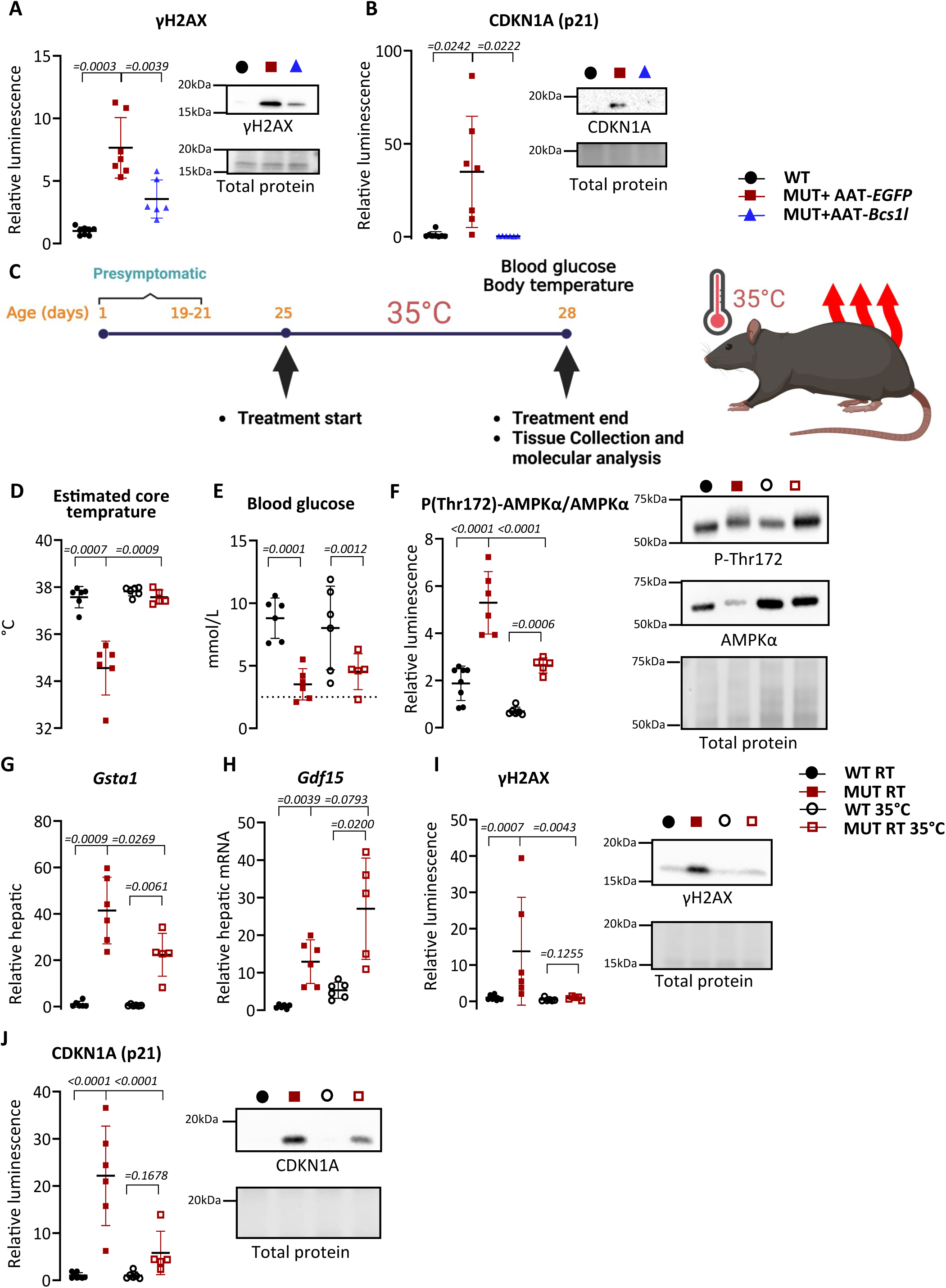
Forced thermoneutrality improves energy homeostasis and prevents cellular senescence. A and B) Western blot quantification and representative blot of γH2AX and CDKN1A (p21) from P28 liver lysates (*n*=5-8/group). C) Schematic presentation of the of the thermoneutrality experiment. D-J) Effect of the elevated housing temperature on D) estimated core temperature (n=5-6/group), E) blood glucose (n=5-6/group), F) hepatic AMPKα phosphorylation (*n*=5-8/group), G and H) hepatic *Gsta1* and *Gdf15* mRNA expression (n=5-6/group), I and J) hepatic γH2AX and CDKN1A levels (*n*=5-8/group). Statistics: one-way ANOVA followed by the selected pairwise comparisons with Welch’s t-statistics (A, B, I & J), Welch’s *t*-test (D & E), and Mann-Whitney U test (G & H). P values of the group comparisons are indicated above the panel. The error bars stand for standard deviation. All data points derive from independent mice.

## Discussion

To probe the systemic roles of the liver in mitochondrial disease pathology, we performed a preclinical rAAV-based intervention in the patient mutation-carrying *Bcs1l^p.S78G^* mouse model of CIII deficiency. We show that a single intraperitonial injection of hepatocyte-specific rAAV-*Bcs1l* into juvenile pre-symptomatic mice prevents liver disease, improves growth, corrects hypothermia, and doubles the survival. The immediate cause of the extended survival was likely the prevention of lethal metabolic crisis, i.e. extreme hypoglycemia. Several liver-intrinsic or -dependent processes may underlie these robust therapeutic effects. The growth from the fetal period until the end of adolescence requires substantial amount of energy and metabolic resources. In GRACILE syndrome patients the growth restriction is of fetal onset^6^, whereas the growth of *Bcs1l^p.S78G^* mice rapidly slows down after 3 weeks of age, coinciding with a linear decrease in CIII activity^8^. Hepatocyte-targeted gene therapy partially restored hepatic *Igf-1* and *Ghr* expression and improved growth, which indicates improved resources for growth and other energy-consuming processes like thermoregulation. The major unexpected findings were the previously unrecognized pathological hypothermia and BAT dysfunction in the *Bcs1l^p.S78G^* mice, as well as normalization of their body temperature solely by the hepatocyte-specific restoration of CIII function. Mitochondrial diseases disrupt energy metabolism, and the largest consumer of energy is basal metabolism^31^. Surprisingly, only a few reports have mentioned hypothermia in patients with mitochondrial disease^39–41^. Perhaps the most interesting of these states that the brain is hypothermic in mitochondrial disease patients^39^. In newborn infants with GRACILE syndrome, hypothermia has not been noted, possibly becauses the babies are cared in incubators with ambient temperature adjusted to provide a normal core temperature.

Basal metabolism, or the resting metabolic rate, accounts for >50% of the total energy expenditure in the fed state at thermoneutrality in mice^42^, and even more at the normal housing temperature. In juvenile *Bcs1l^p.S78G^*mice, CIII activity is broadly decreased, by 50-90%, including in the liver and skeletal muscle, the largest heat-producing organs. Correspondingly, the weight-normalized night-time energy expenditure is 25-50% lower than that of wild-type littermates^11^. The mice are also small, which increases relative heat loss via skin, and they lack dermal adipose tissue^10^. These aspects undoubtedly result in a great strain on the adaptive thermoregulatory system. Adaptive (also called facultative) thermogenesis is carried out by the skeletal muscle and brown and beige adipose tissues, and is under neuronal control^31^. BAT is a major site for heat production in newborn humans and in small homeothermic animals like mice^31^. In a healthy animal, thermoreceptors in the skin and visceral organs monitor temperature and transmit the signals via the hypothalamus to sympathetic nerve termini in the BAT. Surprisingly, ADRB3 signalling and UCP1 expression were unaffected in the hypothermic mutant mice at RT and, importantly, after an acute cold exposure, indicating that their BAT is unresponsive to cold. On the basis of nociception (hot plate) data from adult mutant mice, we first hypothesized that defective thermosensation may underlie the lack of BAT activation. Antibody staining of foot pad tissue sections showed disrupted sensory innervation at the dermis/epidermis boundary. Physiologically, eliminating sympathetic neuronal stimulation from hypothalamus to BAT via surgical denervation indeed leads to dampened UCP1 induction upon feeding (diet-induced thermogenesis). Basal UCP1 expression is not affected^43^, similarly to the *Bcs1l^p.S78G^* mice. Conversely, the expression of all three β-adrenergic receptors (*Adrb1-3*) in BAT is decreased by 50%-60% in *Ucp1* germline KO mice^44^, indicating that defective mitochondrial uncoupling in BAT can lead to downregulation of the β-adrenergic receptors, also a scenario compatible with our findings. The sympathetic innervation of final target tissues commences during late embryogenesis in mice and continues for 3–4 weeks after birth^45^. Therefore, our findings in the 4-week-old mice likely represent a postnatal neurodevelopmental phenotype; to our knowledge the first such reported in any mouse model of primary OXPHOS deficiency. Peripheral neuropathies that are quite common among mitochondrial diseases are typically later childhood- or adult-onset neurodegenerative phenotypes^46^. The reduced hearing sensitivity in adult *Bcs1l^p.S78G^* mice and the sensorineural hearing loss in patients with other *BCS1L* mutations^47^ are consistent with peripheral neuropathy.

Presumably controlled by neuronal adrenergic signaling similar to BAT in rodents^48^, we expected that also the induction of skeletal muscle thermogenesis would be impaired in the mutant mice. However, the mice were apparently able to signal the hypothermia to skeletal muscle to initiate a thermogenic response. Although skeletal muscle thermogenesis indeed seemed to be activated in the CIII-deficient mice, it was clearly insufficient to maintain euthermia, probably due to the low CIII activity (25% of WT at P30^11^) in this tissue, as well as simultaneous low hepatic thermogenesis. In an early whole body calorimetry (CLAMS) experiment, we inadvertently found out that the *Bcs1l^p.S78G^*mice are highly cold-sensitive, when all mutant mice were found dead after a 12-h +4°C cycle in the CLAMS program. The mice behave somewhat similarly to *Ucp1;Sln* double knock-out mice, which are viable but extremely sensitive to cold, with even standard housing temperature causing neonatal lethality^49^. Interestingly, the *Ucp1;Sln* double knock-out mice survive the cold if adjusted gradually, but lose weight and WAT fat stores.

In rodents, BAT starts to develop from E15–16 and continues to develop postnatally for 2-3 weeks^50^. The *Bcs1l^p.S78G^*mice are healthy until 3 weeks of age, making it unlikely that they have a devepomental BAT defect. Instead, our data strongly points to acquired BAT dysfunction caused by the low CIII activity, which is well below the 50% threshold for incipient pathology in other mouse tissues^8,11^. Quite surprisingly, there are no prior studies of thermogenesis or thermoregulation in mitochondrial disease models. However, artificial models with various adipose tissue-specific knock-outs (KOs) of Mitochondrial transcription factor A (TFAM) have been reported and have shown that severe mitochondrial dysfunction can lead to BAT inflammation and dysfunction. For example, adipose tissue (*adiponectin-Cre*) KO of TFAM resulted in broad OXPHOS deficiency in white adipose tissue (WAT) and BAT, BAT inflammation, lipodystrophy, and hepatosteatosis^51^. BAT-specific (*Ucp1-Cre*) KO of Optic atrophy 1 (OPA1) leads to mitochondrial dysfunction in BAT while, counterintuitively, improving the energy balance and thermoregulation of the mice via compensatory browning of WAT^52^. As for the possible compensatory browning, we were unable to study the WAT compartment because, similarly to GRACILE patients, the juvenile *Bcs1l^p.S78G^* have no detectable WAT depots, a phenotype we have attributed to their progeroid disease^10^. Though not a human disease model, the mutator mice, a widely studied mouse model of OXPHOS dysfunction due to a proofreading-mutant mitochondrial DNA polymerase γ, develop hypothermia after about seven months of age^53,54^. The mutator mice show decreased BAT temperature and repressed thermogenic gene expression^55^. Given so few relevant models and studies, there is little understanding of the mechanisms via which mitochondrial dysfunction leads to broader BAT dysfunction beyond defective respiratory electron transfer. Activated BAT uptakes glucose vigorously, which is utilized in positron emission tomography (PET) imaging, and the uptake could be compromised by the hypoglycemia in the *Bcs1l^p.S78G^*mice. However, the main activators and fuel molecules for thermogenesis are fatty acids rather than glucose, which is mostly used for lactate and triglyceride (TG) synthesis in BAT. Fatty acids are simultaneously esterified into TGs and hydrolyzed in brown adipocytes to fuel thermogenesis^56^. *De novo* lipogenesis and esterification consume ATP, which is why ATP synthesis by OXPHOS is crucial to thermogenesis, in addition to maintenance of membrane potential for uncoupling^56^. The detrimental consequences of blocked BAT FAO were demonstrated in mice with adipose-specific knockout of carnitine palmitoyltransferase 2 (CPT2A)^23^. The *Cpt2a^−/−^* mice fail to upregulate the thermogenic genes *Ucp1*, *Pgc1α,* and *Dio2* in response to an ADRB3 agonist or acute cold, and become fatally hypothermic upon acute cold challenge. The data also suggesed that loss of BAT FAO feeds back to inhibit FAO-related gene expression in BAT^23^, a scenario that is plausible in the *Bcs1l^p.S78G^* mice as well. In another striking example, *Atgl* (adipose triglyceride lipase, current gene name *Pnpla2)* knock-out led to BAT whitening (fat accumulation) and inflammatory macrophage infiltration with crown-like macrophage figures apparent in galectin 3 antibody staining^57^. We found that gene expression related to FAO and browning were suppressed in the CIII-deficient BAT whereas inflammatory markers were highly increased. Intriguingly, markers for a newly characterized fibrogenic macrophage population named Fab5 macrophages^25^, were highly upregulated in the liver and BAT of the *Bcs1l^p.S78G^* mice. Galectin-3 immunostaining showed abundant crown-like macrophages in the mutant BAT. In summary, we conclude that compromised OXPHOS and FAO lead to BAT dysfunction and inflammation in the CIII-deficient mice.

The liver is a major metabolic hub with historically known but relatively little studied roles in thermogenesis and thermoregulation^58–62^. First studies indicating the liver as a major thermogenic organ were published decades ago^60,63^, but there is very scarce newer literature on the topic^62^, perhaps because the field has since the 1970s focused on BAT thermogenesis, particularly in the context of obesity^64^. Hepatic heat originates from basal mitochondrial and non-mitochondrial metabolisms, mainly ATP hydrolysis, in which 60% of the energy can be released as heat. Strikingly, AAT-*Bcs1l* prevented the hepatic glycogen depletion, fat accumulation, and normalized fuel utilization-related gene expression despite the continued low CIII activity in many other tissues. The gene expression data also suggested paradoxical downregulation of fatty acid oxidation in the CIII-deficient liver. As we suggested in an early metabolomics study^65^, this could be due to compromised β-oxidation, which feeds electrons to the CoQ pool via electron-transferring flavoprotein dehydrogenase and is thus dependent on CoQ oxidation by CIII^5^. The FAO-related gene expression was fully normalized by the gene therapy, alleviating the energy metabolic stress locally in the liver and systemically. Interestingly, in the older (P150-200) mutants in the longer-living mtDNA background, FAO appear to be eventually upregulated and the livers lack fat accumulation^19,66^, showing eventual adjustment of the energy metabolism to the chronic CIII deficiency. The most straightforward interpretation of our temperature, hepatic gene expression, and histologial data is that normalization of basal liver energy metabolism restored basal metabolic heat production. The liver being a large organ with very high blood flow, this was sufficient to maintain euthermia systemically, even with inactive BAT. Curiously, the fact that AOX-expressing mutant mice maintained near-normal core temperature suggests that prevention of hypotermia was a key factor in its broad beneficial effects in *Bcs1l^p.S78G^* mice, which we have reported earlier^9,10^. Among the effects reversed by AOX, cell proliferation against lack of energy and biosynthetic resources leads to DNA damage, cell cycle arrest and senescence in the *Bcs1l^p.S78G^*mice, contributing to the liver pathology^10^. Here, AAV-mediated restoration of CIII activity completely prevented the regenerative hepatocyte proliferation, redirecting also those resources to systemic energy metabolism and growth. Housing the mice at 35°C, but not at the typical 30°C considered thermoneutral for WT mice, was sufficient to relieve the metabolic stress. This most likely abolished the DNA damage and senescence markers.

Our findings shed light on the intricate interplay between mitochodria, hepatic energy metabolism, and adaptive thermogenesis and thermosensation in thermoregulation. For the first time in a physiologically relevant patient mutation-carrying mouse model, we describe BAT dysfunction caused by primary OXPHOS deficiency. With these findings we hope to reintroduces the largely forgotten critical role of the liver as a thermogenic organ. Finally, our study highlightings the complexity of mitochondrial disease pathophysiology and the potential of tissue-targeted gene therapy to produce robust therapeutic effects despite, or sometimes because of, the complexity.

### Limitations of the study

The translatability of the results to human CIII deficiencies remains to be studied. Thermogenesis and thermoregulation are quite different between human and mice simply because of the body size difference, small animals being much more prone to heat loss. It may be that hypothermia is mostly a mouse model-specific manifestation and rare in patients with mitochondrial diseases, although this should be studied more thoroughly clinically. With the current set of experiments it was not possible to make a firm conclusion about the effect of the defective skin sensory innervation to thermosensation and subsequent induction of adaptive thermogenesis. More detailed analysis of the sympathetic nervous system was out of the scope of this study, but neurophysiological measurements and analyses of the sympathetic ganglia for, for example, temperature-sensitive ion channel expression would be warranted to clarify the scope of the phenotype. The long-term safety and efficacy of rAAV-mediated gene therapy in CIII deficiency patients carrying *BCS1L* mutations and with widely varying phenotypes remain to be established in clinical studies.

## Materials and Methods

### Cloning and virus production

Mouse *Bcs1l* coding sequence (*MmBcs1l*) was PCR-amplified from mouse tissue cDNA (primers EcoRI-MmBcs1l: 5’-ATGAATTCACCATGCCATTTTCAGACTTTGTTCTG-3’ and MmBcs1l-STP-HindIII: 5’-ATAAGCTTTCACCTCAGAGATTCAATGTTGT-3’). The fragment was ligated into pBluescript and sequenced. The insert was subcloned into a pAAV2-LSP1-PB(TR)-EGFP vector, which drives hepatocyte-specific expression under the human ApoE enhancer and α1-antitrypsin (AAT) promoter (generously provided by Professor Ian Alexander, University of Sydney, Australia). The original vector containing Enhanced Green Fluorescent Protein (EGFP) cDNA was used as a control. Another construct was prepared in the same vector backbone containing *AOX* from *Crassostrea gigas* (*CgAOX*). This was done by subcloning from pcDNA6A-*CgAOX* construct^67^ Along with the expression cassette, the vectors also contained flanking PiggyBac transposon recognition sequences, which allow genomic integration upon parallel expression of PiggyBac transposase^16^. Serotype 9 viral particles were produced by the AAV Gene Transfer and Cell Therapy Core Facility of the University of Helsinki.

### Mouse breeding and husbandry

The animal facilities of the University of Helsinki maintained the mice on the C57BL/6JCrl background (Harlan stock 000016). An *Bcs1l^p.S78G^* mutant strain carrying a spontaneous *mt-Cyb^p.D254N^* variant^11^ was used throughout this study, with the exception of the thermoneutrality intervention and hot plate response measurement (see section below). In the mtDNA background, the homozygotes become terminally ill at approximately 1 month of age, allowing fast assessment of survival. *Bcs1l* wildtype or heterozygous animals were used as healthy controls (wild type, WT, by phenotype) controls. *Bcs1l^p.S78^*;*mt-Cyb^p.D254N^* mice carrying a single copy of the *Ciona intestinalis* alternative oxidase (AOX) transgene were also used for this study. Both males and females were used and data are shown separately for them only if a significant sex differences were observed or known.

The mice were housed in individually ventilated cages with a 12-hour light/12-hour dark cycle at a temperature of 22-23°C, and they had ad libitum access to water and food (2018 Teklad global 18% protein rodent diet, Envigo). Mouse health was monitored by manual behavioral scoring and weighing according to the ethical permit. Samples were collected at postnatal day 28 (P28) or according to the survival of the mice. In survival analysis, the mice were euthanized when weight loss was greater than 15% of the maximum weight of the individual mouse.

The animal studies were approved by the animal ethics committee of the State Provincial Office of Southern Finland (ESAVI/16278/2020 and ESAVI/31141/2023) and were performed according to FELASA (Federation of Laboratory Animal Science Associations) guidelines. At the German Mouse Clinic (GMC), the mice were maintained from 8 weeks of age until sacrificing at week 21 according to the GMC housing conditions (www.mouseclinic.de) and German laws.

### rAAV administration

Presymtomatic (postnatal day 19-23, P19-23) mutant mice were injected intraperitoneally with 100 µl saline containing 5×10^10^ viral particles encoding EGFP, wild-type BCS1L or AOX. Following the administration, the health of the mice was monitored visually and by weighing according to the ethical permit.

### Thermoneutrality intervention and warm temperature treatment

A group of *Bcs1l^p.S78G^* mice in WT mtDNA background were housed at 30-32°C for 10 days and sacrificed at P33. Further, a group of *Bcs1l^p.S78G^*; *mt-Cyb^p.D254N^* mice was kept in 35°C for 3 days and sacrificed at P28. Mouse health was monitored by manual behavioral scoring and weighing according to the ethical permit.

### Assessment of nociception

Hotplate test (thermal pain response) weas carried out at the German Mouse Clinic, Munich, Germany on a group of *Bcs1l^p.S78G^* mice in WT mtDNA background of P100^9^.

### Assessment of body composition and temperature

Echo magnetic resonance imaging (Echo-MRI) based MiniSpec Body Composition Analyzer (Bruker, Billerica) was used to quantify fat mass. For measuring skin temperature, Gentle Temp 720 IR meter (Omron Health Care) was used. The core temperature was estimated based on the skin temperature of sternum with the following formula: *Tcore* = −163.9 + 11.47 ∗ *Tskin* − 0.1629 ∗ *Tskin*^2^. This formula derives from in-house calibration data from 33 mice from which both skin temperature and core temperature were measured. The core temperature was measured by directly inserting the thermometer in the abdominal cavity. The estimated core temperature was used when the temperature data from the abdominal cavity was not available. An infrared camera (FLIR C2, Teledyne FLIR LLC) was used to acquire a dorsal thermal image.

### Necropsy

The mice were euthanized by cervical dislocation. Before sample collection, the mice underwent a 2-hour fasting period, and all samples were obtained at similar time of the day, during the light period. Skin temperature was measured before euthanasia. Tissues were either placed in 10% histology-grade formalin or frozen in liquid nitrogen and stored at -80°C. Blood glucose was measured with a stick meter (Freestyle lite, Abbott) from the blood within the body cavity while collecting the other samples.

### Quantitative PCR (qPCR) and RNA sequencing

Total RNA was extracted from the snap-frozen tissue samples with RNAzol RT reagent (Sigma-Aldrich). qPCR was performed from the cDNA using EvaGreen- and Phire II Hot Start DNA polymerase-based detection chemistry^11^. CFX96 thermocycler and CFX Manager software (Bio-Rad) were utilized to perform the qPCR and data analysis. LinRegPCR software^68^ was used to calculate the PCR efficiency. List of all primers used can be found in Supplementary Table 1. *Gak* and *Rab11a* served as reference genes.

RNA sequencing and basic bioinformatics of the data were carried out by Novogene Inc. (service: Plant and Animal Eukaryotic mRNA-seq with reference, WBI-Quantification), using poly-A enriched libraries and NovaSeq X Plus Series (PE150) platform. 9 Gb of raw data per sample were obtained.

### SDS-PAGE, blue-native PAGE and Western blot

For Western blot analyses of tissue samples, 5-10 mg frozen tissue samples were homogenized in SDS lysis buffer (2% SDS, 60 mM Tris-Cl pH 6.8, 12.5% glycerol, 1 mM EDTA) supplemented with protease inhibitors (Complete protease inhibitor mix, Roche). Sample viscosity was reduced by sonication. To determine protein concentrations, Bradford reagent (Bio-Rad) was employed, and bovine serum albumin standards were used as reference. To mitigate interference from SDS, 2.5 mg/ml of α-cyclodextrin (Sigma-Aldrich) was added to the reagent^69^. For regular SDS-PAGE, bromphenol blue and 5% β-mercaptoethanol were added to the protein lysates. The samples were heated at 95°C for 5 minutes, and 5–30 µg total protein per lane run in commercial gradient gels (Bio-Rad). Transfer to polyvinylidene fluoride (PVDF) (Amersham Hybond P 0.2 PVDF, Cytiva) membrane was achieved through the tank transfer method using standard Towbin transfer buffer containing 10% MeOH. To verify equal loading and transfer, the membranes were stained with Coomassie G-250. A list of all antibodies used can be found in Supplementary Table 2. Peroxidase-conjugated secondary antibodies, enhanced chemiluminescence, and the Bio-Rad ChemiDoc MP Imaging System with Image Lab software (Bio-Rad) and LI-COR Odyssey M Imaging System and acquisition sotware were used for the detection and documentation of immunoreactions. All the samples were randomized before running and processing for quantification. For representative blots, individual samples were pooled from each experimental group to obtain an average signal, which represents each group.

For blue native gel electrophoresis (BNGE), liver mitochondria were first freshly isolated using the previously described protocol^11^. Then they were solubilized by adding 6 mg digitonin per mg of protein in cold buffer comprising (50 mM Bis-Tris-Cl^+^, 50 mM NaCl, 1.4mg/ml digitonin, 10% glycerol, 1 mM EDTA, protease inhibitor mix, pH of 7.0). The sample lysates were clarified by centrifugation at 18,000g at +4°C for 6 minutes. Coomassie Blue G-250 (1.5 mg/ml) was added to the supernatants and 10 μg solubilized mitochondrial protein separated using 3–12% NativePAGE^TM^ Bis-Tris gradient gels (Invitrogen) and electroransferred onto PVDF filters as described^22^. Preparation of RNA library, transcriptome sequencing gene expression change analysis was conducted by Novogene Co., LTD.

### Assessment of respiratory chain enzymatic activities and quantification of ATP

CIII activity was measured from isolated mitochondria using a spectrophotometric method based on 550nm absorbance change upon cytochrome c reduction with decylubiquinol serving as the electron donor^11^. The background was determined in the presence of antimycin A and myxothiazol. The data were normalized to protein content in liver and to complex IV (CIV) activity in BAT.

Enzymatic quantification of ATP from liver was performed using previously described protocol^70^.

### Tissue histology, immunohistochemistry, and immunofluorescence

Formalin-fixed paraffin-embedded tissues underwent standard procedures for general histological assesment, including hematoxylin and eosin (H&E) staining (Sigma-Aldrich) and glycogen detection using periodic acid-Schiff (PAS) staining (Sigma-Aldrich). Additionally, frozen liver and sections fixed in formalin and saturated with 30%_w/v_ sucrose were subjected to the standard Oil-Red-O (ORO) staining (Sigma-Aldrich) to detect triglycerides. The antigen retrieval of paraffin sections for cleaved caspase 3 and HSP60 staining (see supplementary Table 2) was done by immersing the slides in 10 mM Tris-Cl, pH 9.0; 1mM EDTA, and boiled for 15 minutes. After incubating the sections in the primary antibody, ImmPress peroxidase- or alkaline phosphatase polymer-conjugated of secondary antibodies (Vector Laboratories Inc) were added. Nitroblue teterazolium was used to visualize alkaline phosphatase and diaminobenzidine peroxidase activity, respectively. Nuclear Fast Red (Sigma-Aldrich) and hematoxylin were used as nuclear counterstains for alkaline phosphatase and peroxidase stainings, respectively. Image acquisition was performed with Zeiss Axio Imager M2 microscope, Axiocam 503 camera, and Zen 2.3 software (Zeiss).

Sensory neurons were detected by immunofluorescence staining of the hind limb paw foot pads (see supplementary Table 2), followed by counterstaining with Hoechst nuclear stain. Image acquisition was then performed by the Andor Dragonfly spinning disc microscope with Zyla 4.2 sCMOS camera (Andor technology).

### Statistics

All the samples were randomized before running and processing for quantification. For data that followed normal distribution, we applied a one-way ANOVA, followed by selected comparisons using t-tests with Welch’s correction. Where more appropriate, Mann-Whitney U tests were used instead. Survival curves were analyzed using the log-rank Mantel-Cox test. We conducted these statistical analyses using GraphPad Prism 10 software (GraphPad Software Inc.).

## Acknowledgements

We thank Vilma Wanne for technical assistance, professor Ian Alexander for (University of Sydney) for providing the pAAV-LSP1 plasmids. We thank German mouse clinic (Munich) for mouse phenotyping report. We also thank the core facilities of University of Helsinki, FIMM Digital Microscopy and Molecular Pathology Unit, and the Finnish Centre for Laboratory Animal Pathology (Faculty of Veterinary Medicine) for processing of histological samples, Biomedicum Imaging Unit for microscopy services and the Laboratory Animal Center of the University of Helsinki for the animal husbandry. We acknowledge the funding from Folkhälsan Research Center, Jane and Aatos Erkko Foundation, the Foundation for Pediatric Research, Finska Läkaresällskapet, The Liv och Hälsa Foundation, Magnus Ehrnrooth Foundation, the Finnish Society of Sciences and Letters and Finnish Doctoral Programme in Oral Sciences (FINDOS). Graphical illustrations in graphical abstract, Fig. 1A, B, 4K, 6I and 7C were created with BioRender (https://www.biorender.com/).

## Author contributions

R.B., J.P., V.F., S.K. and J.K. designed the study. R.B. wrote the first manuscript draft and prepared the figure panels. R.B., J.P., C.K., and J.K performed the animal experiments and sample collection. R.B, J.P. and J.K. performed the histological analyses. The contributions for other methods were following: body composition analyses (R.B. and J.P.), SDS-PAGE and Western blot analyses (R.B., D.U. and J.K.), Blue-Native PAGE (R.B. and J.P.), qPCR (R.B.), immunohistochemistry (J.K.), ATP measurements (R.B. and J.P) and immunofluorescence staining (T.Z.). R.B. was responsible for the statistics. All authors critically read and commented the manuscript, and R.B. and J.K. revised it accordingly.

## Figure legends

**Supplementary figure 1.**
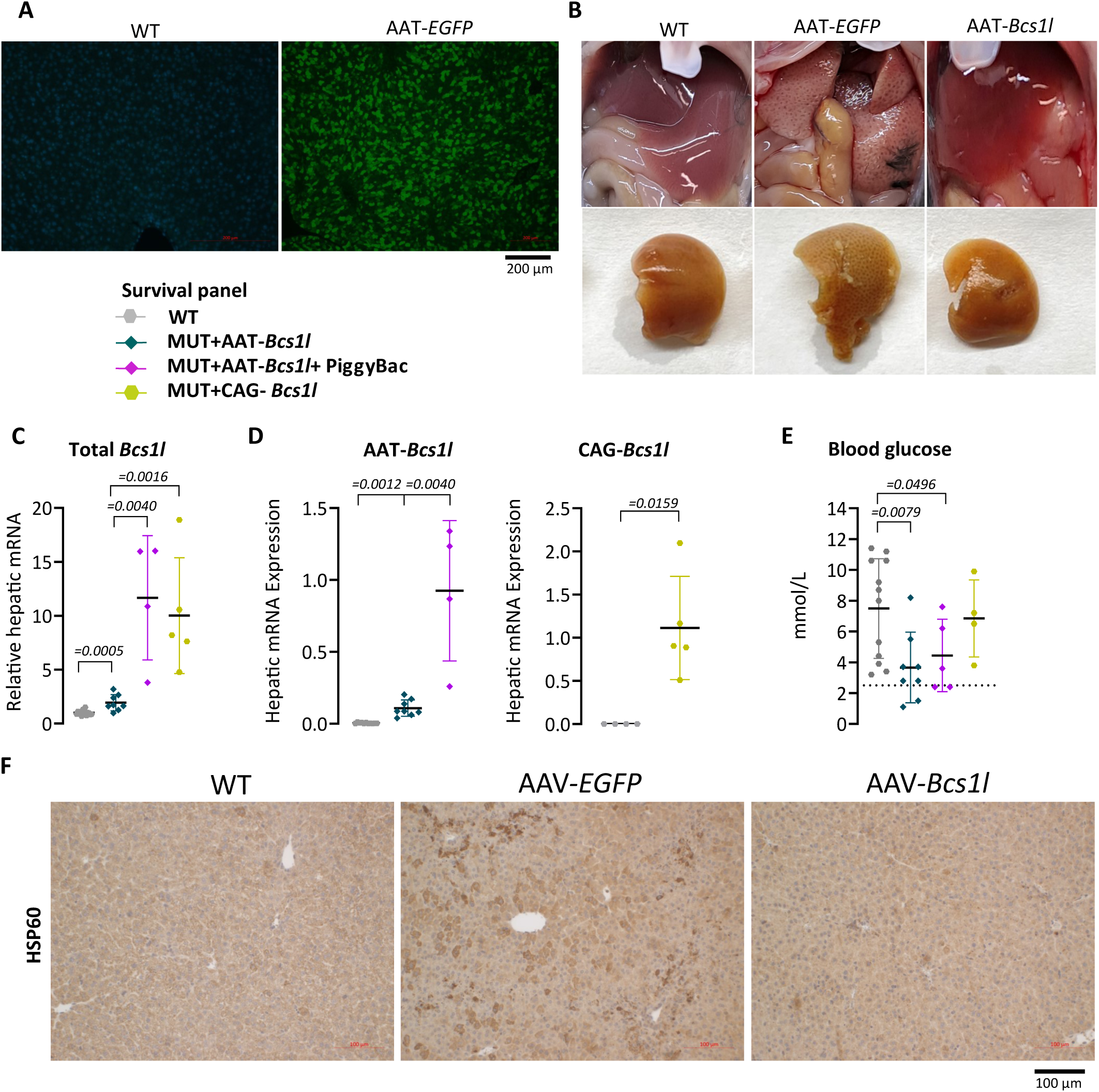
Transduction efficiency of the liver by rAAVs. A) Representative image of transduction frequency with the hepatocyte-targeted viral vector. B) Appearance of the liver at P28 after transduction with rAAV. C) Total *Bcs1l* mRNA at the end stage of the survival analysis (*n*=4-16/group). D) Virally expressed *Bcs1l* mRNA at the end stage of the survival analysis (*n*=4-17/group). E) Blood glucose levels (*n*=4-13/group). The dotted line represents the critical level of glucose (<2.5mmol/L). F) Representative images of liver sections immunostained for the mitochondrial marker, HSP60. Statistics: Mann-Whitney U test (C & D) and one-way ANOVA followed by the selected pairwise comparisons with Welch’s t-statistics (E). P values of the group comparisons are indicated above the panel. The error bars stand for standard deviation. All data points derive from independent mice.

**Supplementary figure 2.**
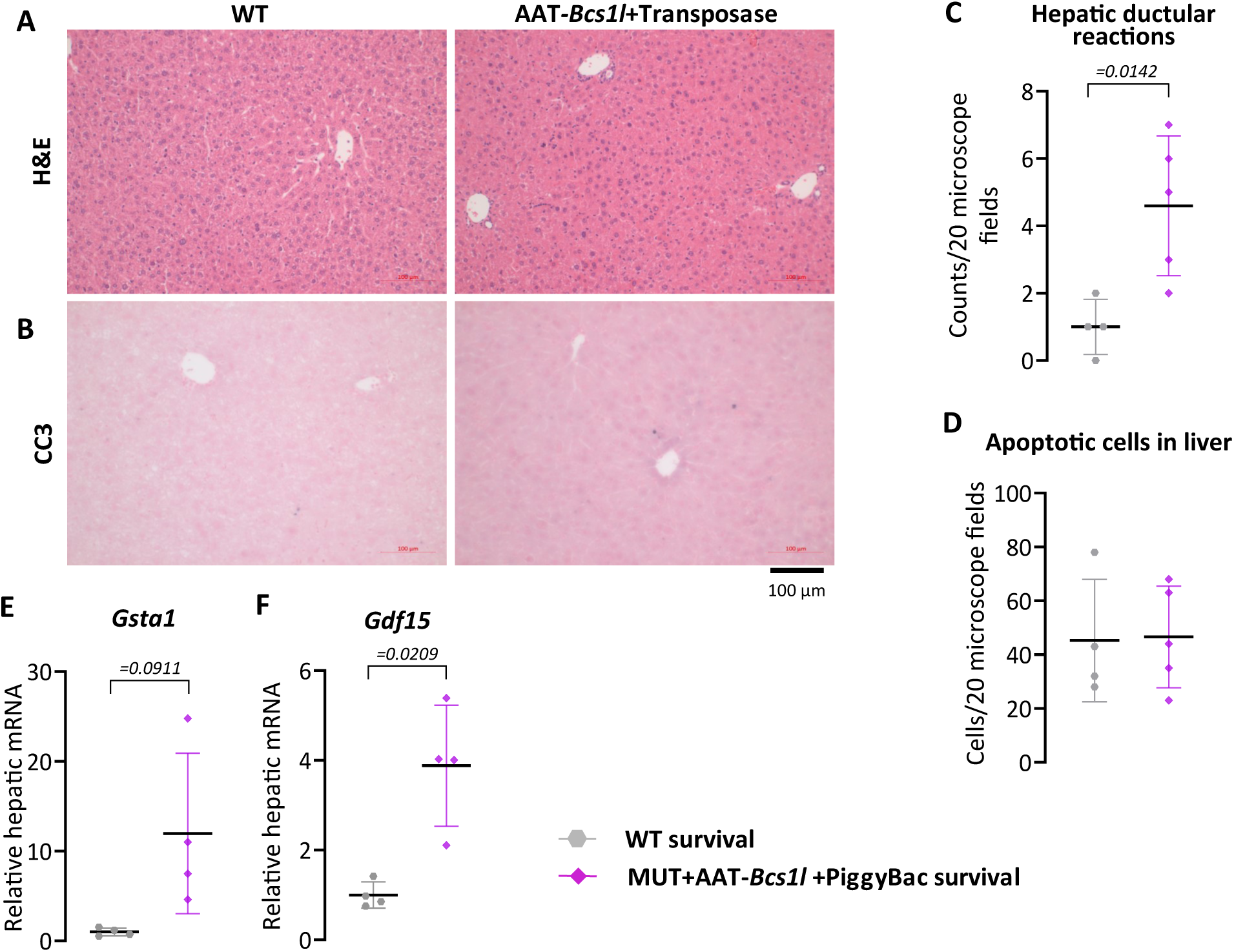
End-stage liver in mutant mice after hepatocyte-targeted gene. A-B) Representative images of H&E-stained liver sections (*n*=5/group) showing tissue. morphology and immunostainings of liver sections for apoptotic cell marker, cleaved caspase 3 (cc3) (*n*=5/group) at the end stage of the survival analysis. C and D) Quantification of the hepatic ductular reactions and the apoptotic cells from H&E-stained sections and cc3-immunostained sections, respectively. E and F) *Gsta1* and *Gdf15* mRNA expression from livers of the survival groups (*n*=4/group). Statistics: Welch’s *t*-test. The error bars stand for standard deviation. P values of the group comparisons are indicated above the panel. All data points derive from independent mice.

**Supplementary figure 3.**
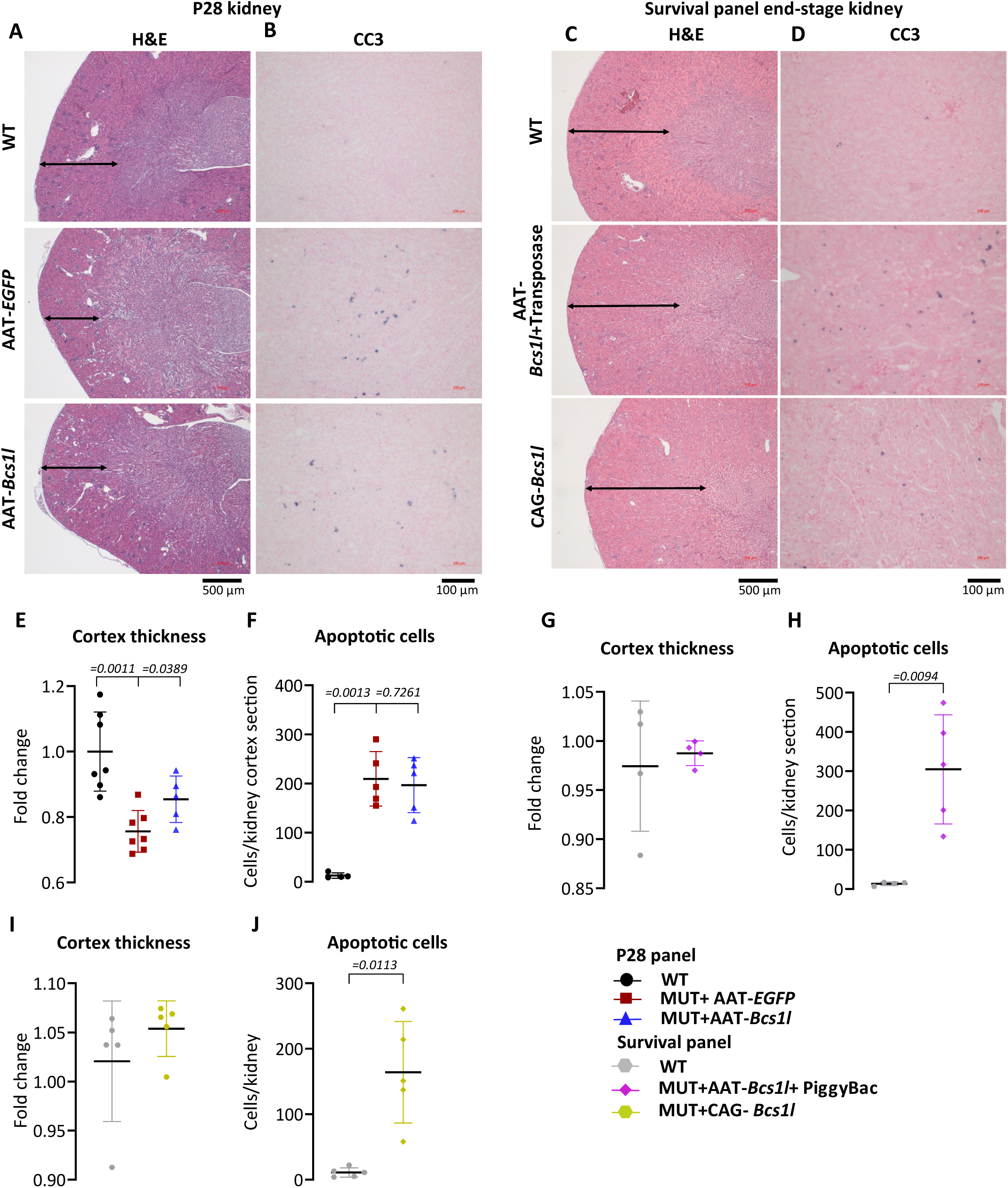
Effect of hepatocyte-targeted gene replacement on kidney disease manifestations. A-D) Representative images of H&E-stained kidney sections (n=4-7/group) showing tissue morphology and immunostainings of cleaved caspase 3 (cc3) (n=5/group) showing apoptotic cells in kidney from P28 mice. E-J) Quantification of kidney cortex thickness and apoptotic cells from H&E-stained and cc3-immunostained sections, respectively from P28 mice, survival AAT-*Bcs1l* +PiggyBac- and CAG-*Bcs1l*-treated mice Statistics: one-way ANOVA followed by the selected pairwise comparisons with Welch’s t-statistics (E and F) and Welch’s *t*-test (G-J). P values of the group comparisons are indicated above the panel. The error bars stand for standard deviation. All data points derive from independent mice.

**Supplementary figure 4.**
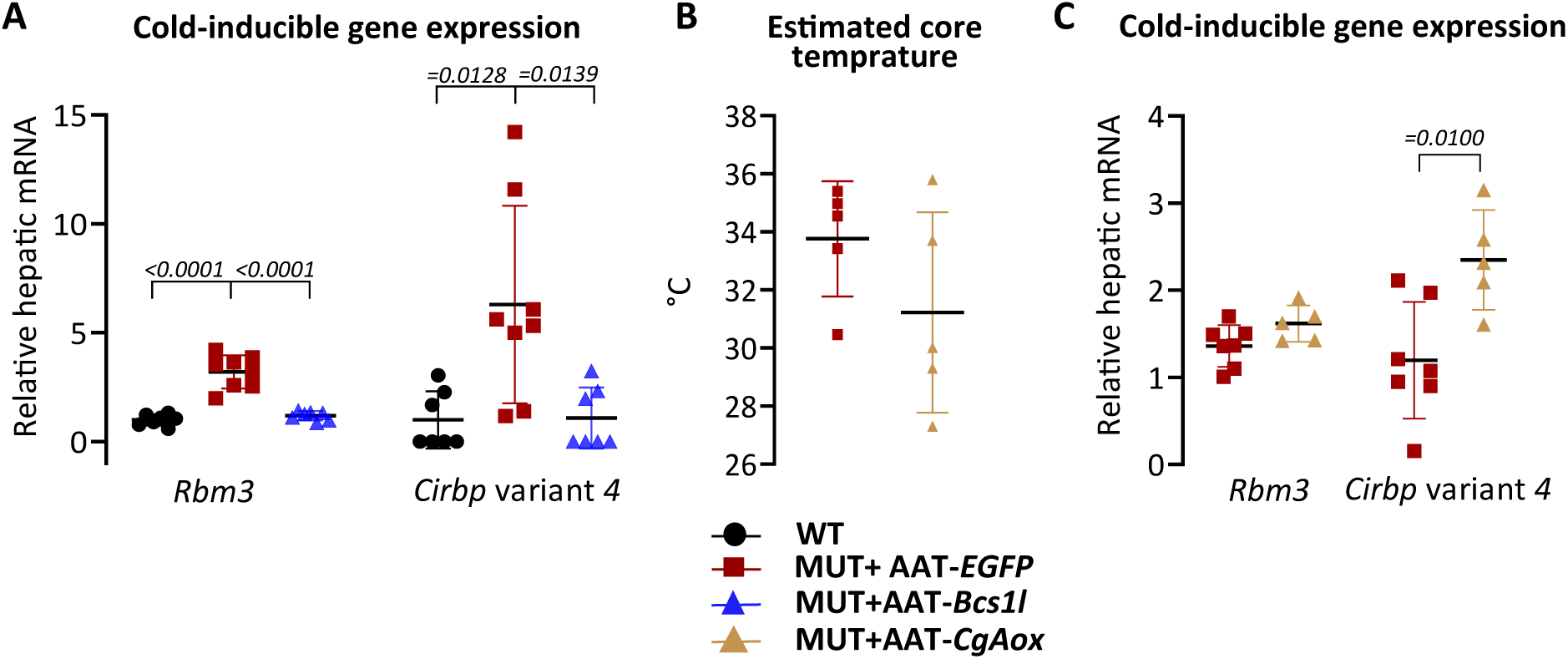
Cold-inducible mRNA expression in CIII-deficient mice. A) Cold-inducible gene expressions from P28 liver (n=7-8/group). B) Effect of *CgAOX* on core temperature at P28 (n=5/group). C) Effect of *CgAOX* on cold-inducible gene expressions in P28 liver (n=5-7/group). Statistics: one-way ANOVA followed by the selected pairwise comparisons with Welch’s t-statistics (A), Welch’s *t*-test (B & C). The error bars stand for standard deviation. All data points derive from independent mice.

**Supplementary figure 5.**
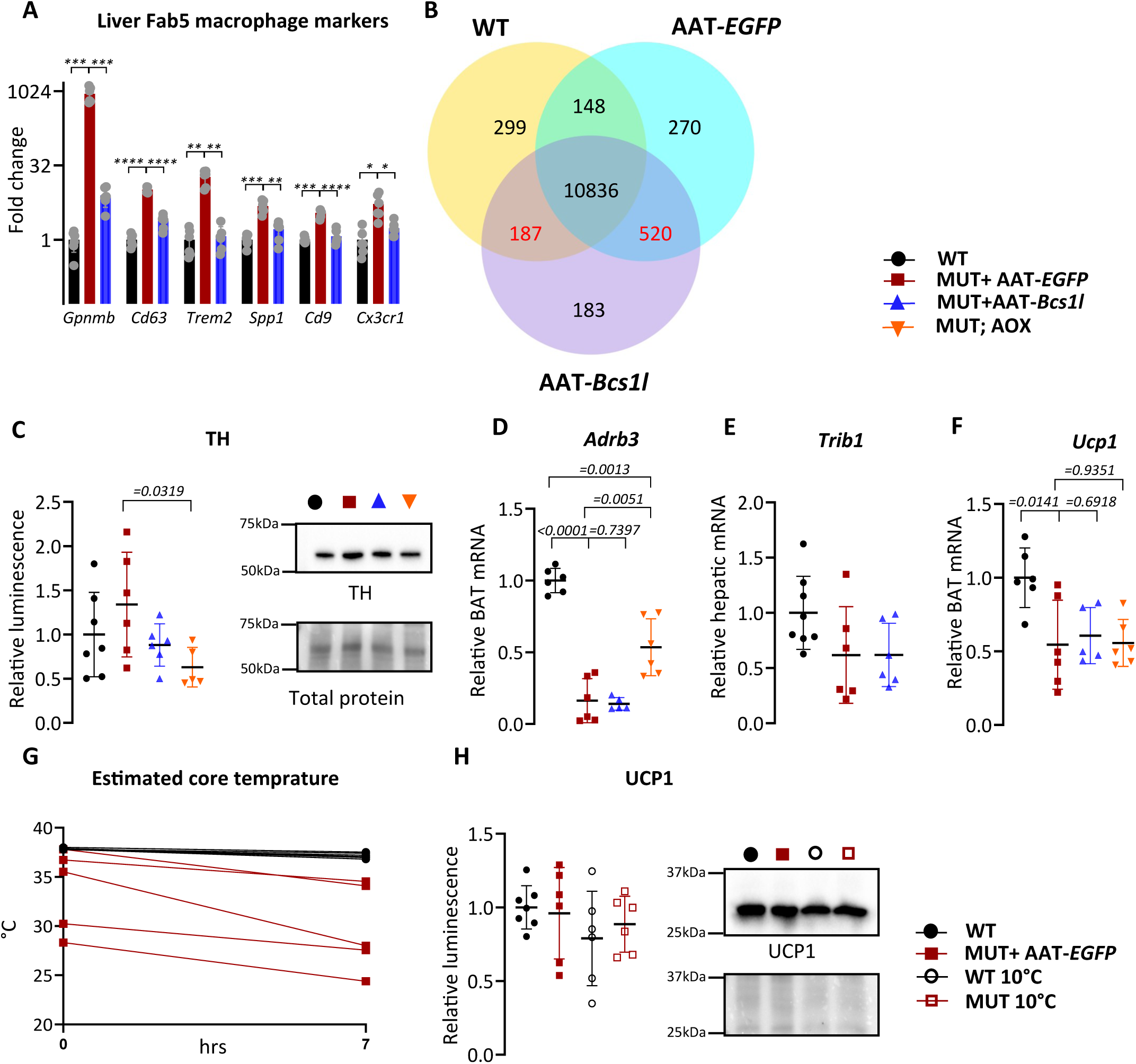
Effects of liver-targeted gene therapy and acute cold exposure on CIII-deficient BAT. A) Fab5 macrophage gene expression in liver transcriptome. B) Differentially expressed genes in BAT transcriptome. C) Western blot quantification and representative blot of tyrosine hydroxylase (TH) from P28 BAT lysates (*n*=6/group). D-F) *Adrb3, Trib1* and *Ucp1* mRNA expression from P28 BAT (n=6/group). A) G) Estimated core temperature before and after 7-hour exposure to 10°C ambient temperature in P28 mice B) H) Western blot quantification and representative blot of uncoupling protein 1 (UCP1) from P28 Statistics: One-way ANOVA followed by the selected pairwise comparisons with Welch’s t-statistics (A, C-F & H). *\*\*\*\*p ≤ 0.0001, ***p ≤ 0.001, **p ≤ 0.01, *p ≤ 0.05* The error bars stand for standard deviation. All data points derive from independent mice.

**Supplementary figure 6.**
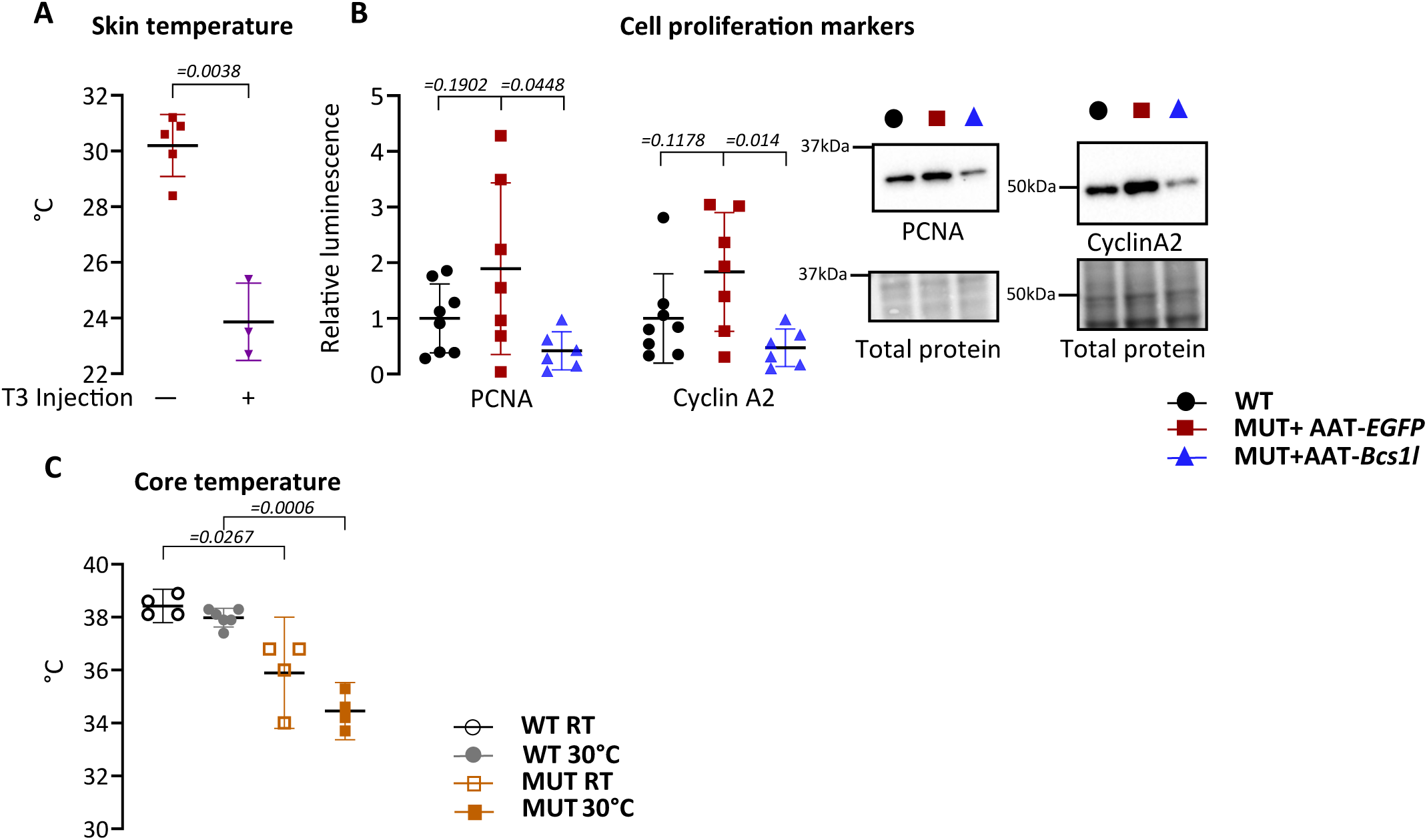
Disease parameters in T3 and AAT-*Bcs1l* injected, and housed in thermoneutral temperature. A) Skin temperature of T3 injected and non-injected P28 mutant mice (n=3-5/group). B) Western blot quantification and representative blot of PCNA and Cyclin A2 from P28 liver lysates (*n*=5-8/group). C) Core temperature of *Bcs1l^p.S78G^* mice at P30 after housing them at (30-32°C) for 10 days (*n*=4-6/group). Statistics: Welch’s *t*-test (A) and one-way ANOVA followed by the selected pairwise comparisons with Welch’s t-statistics (B & C). The error bars stand for standard deviation. All data points derive from independent mice.

**Supplementary Table 1:**
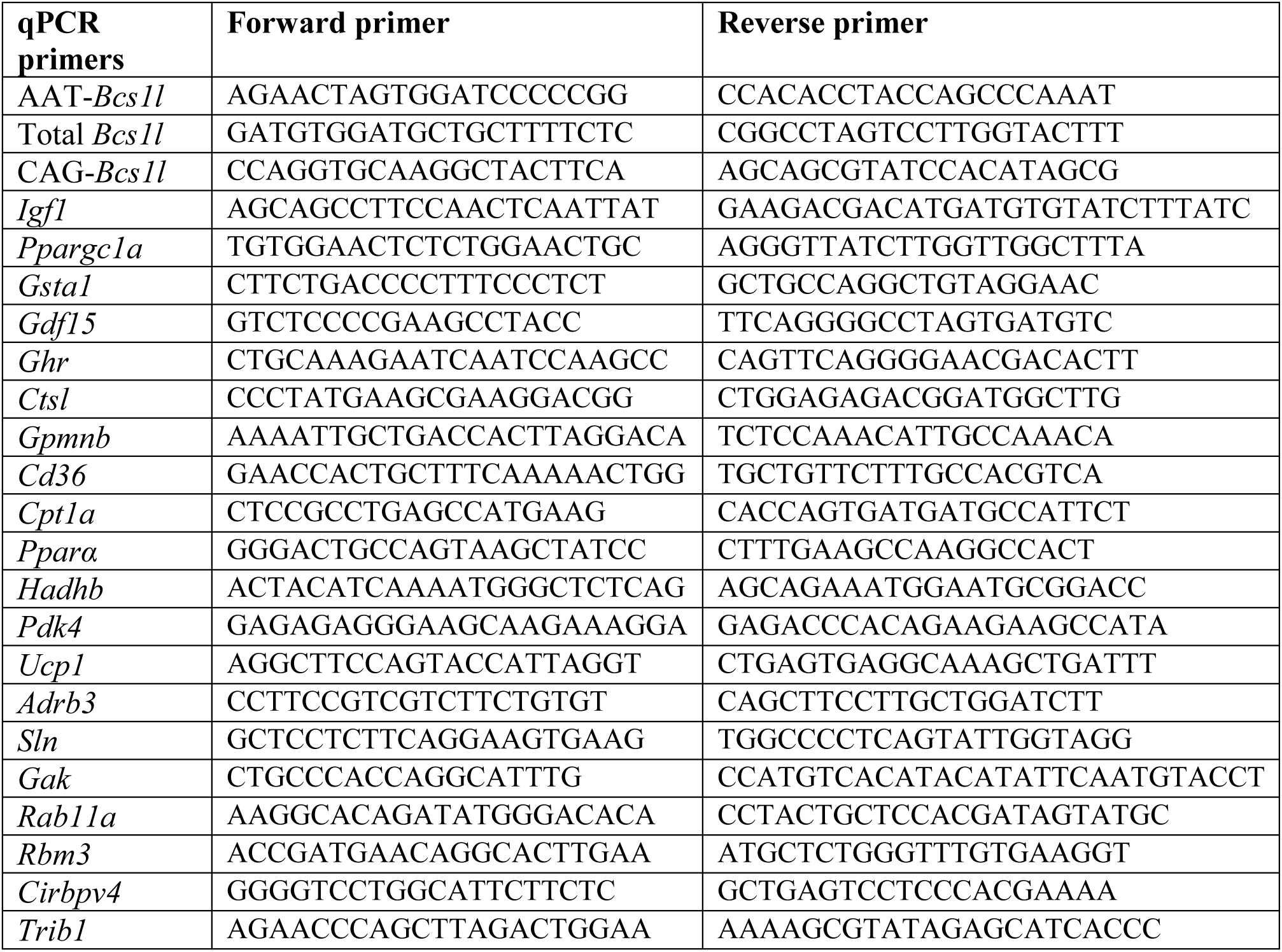
Oligonucleotide sequences in 5’ to 3’ direction.

**Supplementary Table 2:**
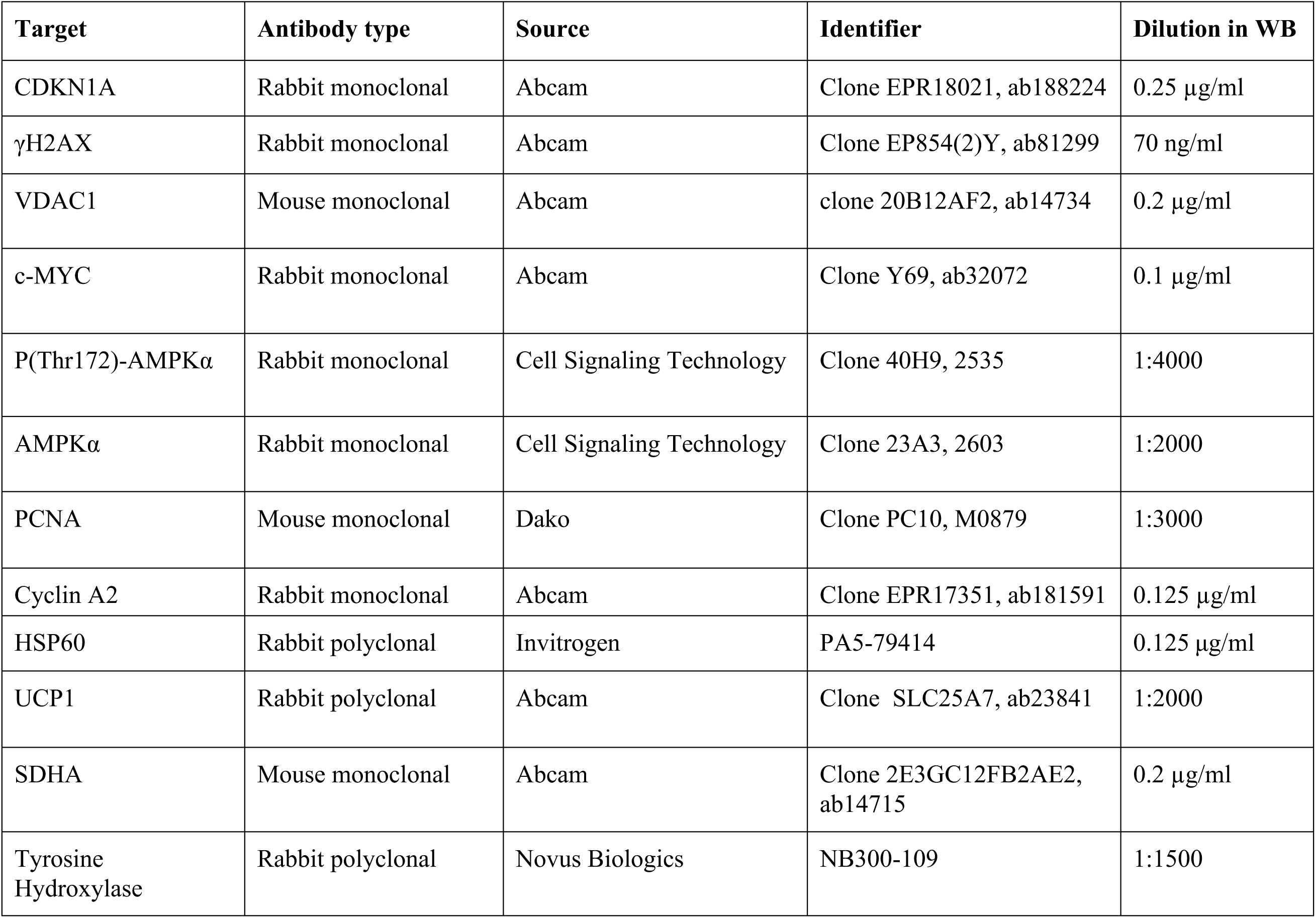
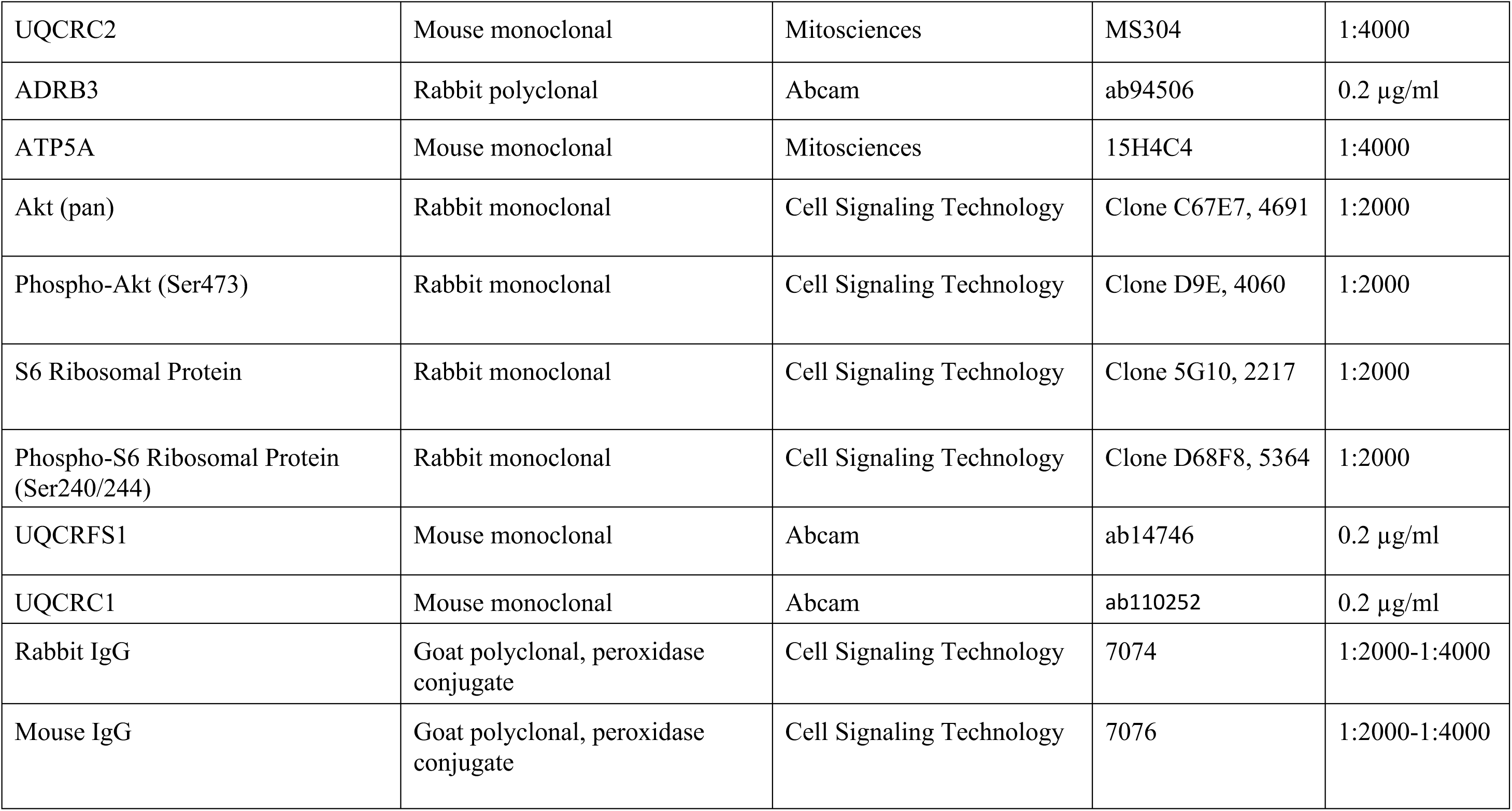
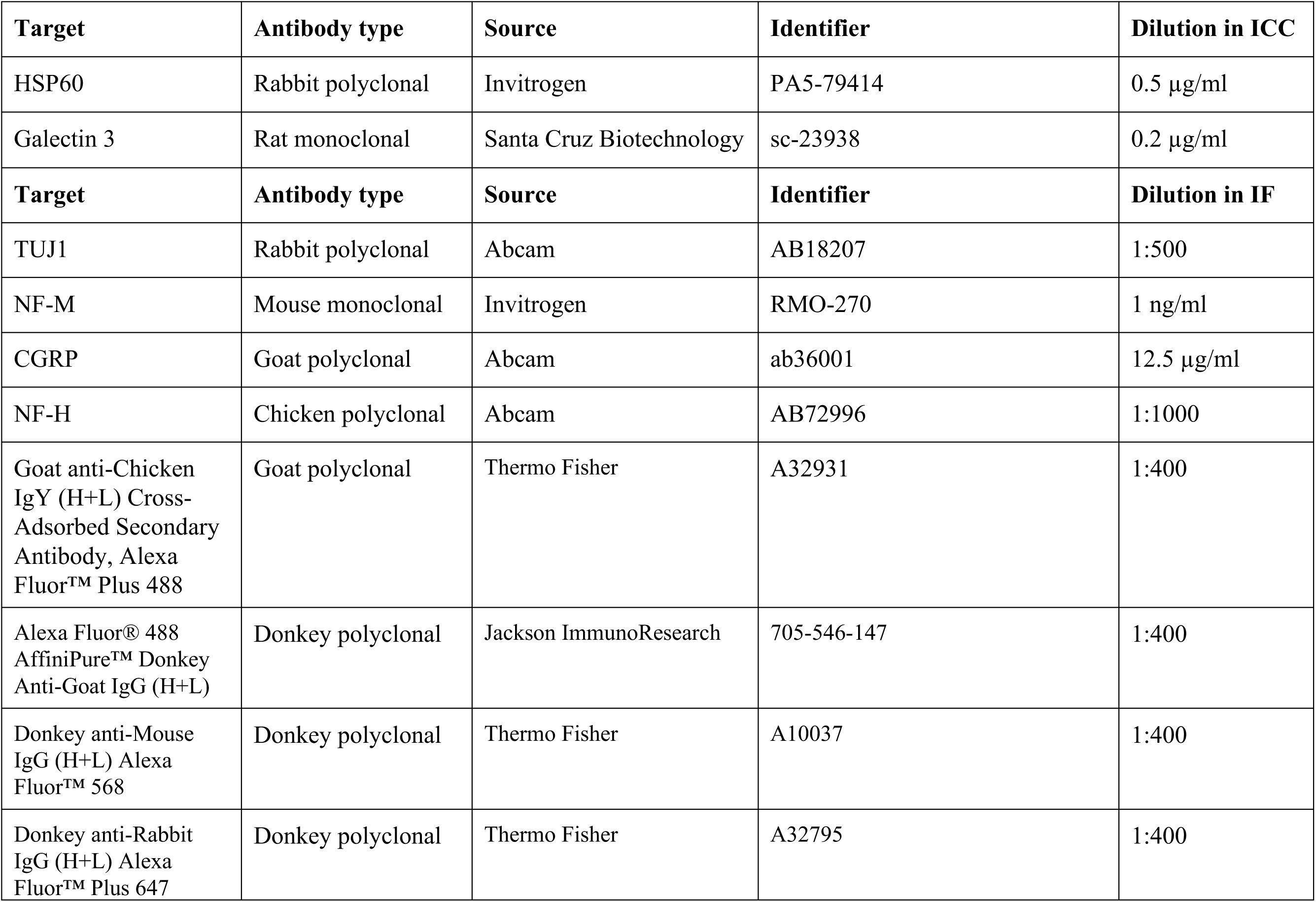
Antibodies.

## Notes

### Competing Interest Statement

The authors have declared no competing interest.

### Summary of Updates

New mouse interventions to assess thermosensation and thermogenesis were performed and the corresponding data added. Transcriptomics data were added and further mRNA and protein data to assess mechanisms. The layout of figure panels and text throughout were revised and amended accordingly. One new author (CK) was added to reflects the additional mouse work.

